# *Paracoccidioides* genomes reflect high levels of species divergence and little interspecific gene flow

**DOI:** 10.1101/2020.07.16.204636

**Authors:** Heidi Mavengere, Katlheen Mattox, Marcus M Teixeira, Victoria E. Sepúlveda, Oscar M. Gomez, Orville Hernandez, Juan McEwen, Daniel R. Matute

**Author notes:** Correspondence: Biology Department, University of North Carolina, Chapel Hill, North Carolina, 250 Bell Tower Drive, Genome Sciences Building, Chapel Hill, NC, 27514, USA.

## Abstract

The fungus *Paracoccidioides* spp. is a prevalent human pathogen endemic to South America. The genus is composed of five species. In this report, we use 37 whole genome sequences to study the allocation of genetic variation in *Paracoccidioides*. We tested three genome-wide predictions of advanced speciation, namely, that all species should be reciprocally monophyletic, that species pairs should be highly differentiated along the whole genome, and that there should be low rates of interspecific gene exchange. We find support for these three hypotheses. Species pairs with older divergences show no evidence of gene exchange, while more recently diverged species pairs show evidence of modest rates of introgression. Our results indicate that as divergence progresses, species boundaries become less porous among *Paracoccidioides* species. Our results suggest that species in *Paracoccidioides* are at different stages along the divergence continuum.

**IMPORTANCE:** *Paracoccidioides* is the causal agent of the most frequent systemic mycosis in Latin America. Most of the inference of the evolutionary history of *Paracoccidioides* has used only a handful of molecular markers. In this report, we evaluate the extent of genome divergence among *Paracoccidioides* species and study the possibility of interspecific gene exchange. We find that all species are highly differentiated. We also find that the amount of gene flow between species is low and in some cases even completely absent in spite of geographic overlap. Our study constitutes a systematic effort to identify species boundaries in fungal pathogens, and determine the extent of gene exchange among fungal species.

## INTRODUCTION

*Paracoccidioides*, a genus of temperature-dimorphic fungi, causes paracoccidioidomycosis (PCM) is a systemic endemic mycosis that occurs across in most countries of Latin America from mid-eastern Mexico to Argentina (Brummer et al. 1993; Restrepo and Tobón 2010; Restrepo et al. 2015). Multiple genetic surveys have revealed extensive genetic variability within *Paracoccidioides* (Morais et al. 2000; Niño-Vega et al. 2000; Feitosa et al. 2003; Kurokawa et al. 2005; Roberto et al. 2016). This variation coupled with the extensive geographic range of the fungus—and the disease it causes—led to the hypothesis of population structure and cryptic speciation within the group. Initial studies reveal the existence of at least three species (Matute et al. 2006) but more recent analyses have suggested the existence of five different species of *Paracoccidioides* (Teixeira et al. 2014; Teixeira et al. 2015; Turissini et al. 2017). Clearly, the use of genetic markers holds the potential to reveal key aspects of the evolutionary biology of the pathogen. Yet, the genetic characterization of most isolates has been modest.

Species from the genus *Paracoccidioides* show a range of divergence that make the group promising to understand how diversification occurs in pathogenic fungi. One of the species of *Paracoccidioides, P. lutzii*, seems to have diverged from the rest of the species at least 30 million years ago (Teixeira et al. 2009). The species show extensive differences in terms of morphology and physiology. Five more species, all within the *brasiliensis* complex form a monophyletic group. *P. restrepiensis*, and *P. venezuelensis* are the most closely related dyad with recent divergence (less than 0.2 MYA; (Turissini et al. 2017a)). *P. brasiliensis sensu stricto* is sister to the *restrepiensis/venezuelensis* dyad while *americana* is sister to the ingroup. *P. brasiliensis sensu stricto* has also been proposed to be formed by two cryptic species: S1a and S1b (Muñoz JF, Farrer RA et al. 2016), but no formal test of this divergence has been performed. All recognized species from the *brasiliensis* complex differ in their morphology. A combination of yeast and conidial morphology differentiates between all species pairs (Teixeira et al. 2015; Siqueira et al. 2016; Turissini et al. 2017)

Whole genome sequences can be used to identify species boundaries in fungi. Three tests jointly indicate the existence of species boundaries (Matute and Sepúlveda 2019). First, genome variation must reflect genetic differentiation in cases where speciation has taken place. In cases of advanced divergence, genomes of putative species should show reciprocal monophyly; this can be measured as the proportion of loci that shows a phylogenetic history concordant with the hypothesized species history. Second, genetic variation should be partitioned across putative species; the extent of genetic differentiation between individuals from different putative species should be larger than the differentiation between individuals within each of the putative species.

Finally, the genomes of putative species should show low or moderate levels of gene exchange. So far, all studies on species boundaries in the genus *Paracoccidioides* have focused on detecting genealogical congruence among a modest number of gene genealogies (Matute et al. 2006; Teixeira et al. 2009; Teixeira et al. 2015; Turissini et al. 2017). Incorporating genomic information is sorely needed to understand the magnitude of differentiation among *Paracoccidioides* pathogens. In this report we use phylogenetics, and population genetics to bridge that gap. We find that the species of *Paracoccidioides* are extensively differentiated which suggest an advanced stage along the divergence continuum.

## METHODS

### Public data

All the data used here has been previously published. The SRA numbers are listed in Table S1. To root our trees (see below), we obtained sequencing reads from two species of *Histoplasma*: *H. capsulatum sensu stricto* and *H. mississippiense* (SRA: Bioproject: PRJNA416769; (Sepúlveda et al. 2017)) These species are among some of the closest relatives of *Paracoccidioides* (Dukik et al. 2017).

### Read mapping and variant calling

Reads were mapped to the *Paracoccidioides brasiliensis* strain Pb18 genome (BioProject: PRJNA28733, BioSample: SAMN02953720), currently assembled into 57 supercontigs, using bwa version 0.7.12 (Li 2013). bam files were then merged using *Samtools* version 0.1.19. Indels were identified and reads locally remapped in the merged bam files using the GATK version 3.2-2 RealignerTargetCreator and IndelRealigner functions (McKenna et al. 2010; DePristo et al. 2011). Subsequently, SNPs were called using the *GATK UnifiedGenotyper* function with the parameter ‘het’ set to 0.01, and all others left as default. The following filters were applied to the resulting vcf file: QD = 2.0, FS_filter = 60.0, MQ_filter = 30.0, MQ_Rank_Sum_filter = - 12.5, and Read_Pos_Rank_Sum_filter = −8.0. Sites were excluded if the coverage was less than 5 or greater than the 99^th^ quantile of the genomic coverage distribution for the given line or if the SNP failed to pass one of the *GATK* filters.

### Ploidy estimation

To detect admixture between species of *Paracoccidioides*, we used *Int-HMM* (Turissini and Matute 2017), an algorithm to detect introgression that requires information on the ploidy of an individual (i.e., it can be run to detect introgression in diploid or haploid organisms; see below). We used genome wide data to determine the most likely ploidy of the *Paracoccidioides* isolates. We used Illumina short reads from the five species of *Paracoccidioides* (described above) to do two ploidy tests. First, we plotted the per site sequencing coverage. In cases in which there is partial aneuploidy in the form of chromosomal duplications, there will be a bimodal distribution. In cases where the genome does not harbor aneuploid regions, there will be a single mode in the distribution. To compare the observed distribution of per site coverage with the null hypothesis of uniform sequencing coverage, we used a Two-Sample Fisher-Pitman Permutation Test (function *oneway_test*, library *coin*; (Hothorn et al. 2008)). We used the ‘*hist’* function (library *graphics*, (R Core Team 2016)) in R to plot the distribution of the per site coverage and of allele frequencies per site across the whole genome for each strain.

Next, we studied the ploidy of *Paracoccidioides* at a local level. We used the same two metrics described above. Sites with the ploidy of the rest of the genome, should show a mean per window coverage. Once-duplicated segments (either as copy number variation or as changes in ploidy) should have twice the coverage that the genome average. We thus calculated the coverage and mean minor allele frequency for each 5kb window in the genome to assess whether there were segments of the genome with evidence for changes in ploidy.

### Phylogenetic reconstructions

Our goal was to determine whether the species from *Paracoccidioides* were reciprocally monophyletic and thus satisfy the requirements to be considered phylogenetic species. We followed a phylogenetic species concept (Taylor et al. 2000; Queiroz and De Queiroz 2007; Matute and Sepúlveda 2019) to recognize species, defining species as genetic clusters that are reciprocally monophyletic and for which there was genealogical concordance across genome-wide unlinked loci. We used two parallel approaches: (*i*) maximum likelihood trees at the genome and at the supercontig level, and (*ii*) Bayesian concordance analysis of the genealogies of orthologous genes. We described each of these two approaches as follows.

#### Maximum likelihood phylogenetics

To determine whether the proposed *Paracoccidioides* species were monophyletic, we first used maximum likelihood phylogenetics. Reciprocal monophyly is a trademark of speciation (Hendry et al. 2009; Matute and Sepúlveda 2019); as divergence accrues, the likelihood of reciprocal monophyly across the whole genome increases for two reasons. First, as divergence increases, the likelihood of incomplete lineage sorting decreases (Gao et al. 2015; Joly et al. 2015). Likewise, in diverged lineages, the magnitude of retained introgression is lower even in cases where hybridization might occur frequently (Muirhead and Presgraves 2016; Turissini and Matute 2017; Hamlin et al. 2020). Since recombination in *Paracoccidioides* occurs but seems to be rare, and there is a high level of linkage disequilibrium across the genome (see *Results*), we studied each supercontig as an unlinked loci. Since mtDNA shows evidence of interspecific gene transfer (Salgado-Salazar et al. 2010; Turissini et al. 2017), we focused only on nuclear genomes We obtained whole supercontig sequences for each individual from the .VCF file using the FastaAlternateReferenceMaker tool in GATK, re-aligned them using Mafft v. 7 (Katoh and Standley 2013), and used them to build Maximum Likelihood (ML) trees using RAxML version 8.2.9 (Stamatakis 2006). We inferred individual trees for each of the largest six supercontigs, which encompass 62% of the genome. We also generated a “genome-wide” tree of a concatenated alignment of all supercontigs. Analyses were run under the GTR+Γ model, with 1,000 bootstrap pseudoreplicates to assess support for each node. The “genome-wide” analysis was partitioned by supercontig, with each partition having its own set of GTR+Γ model parameters, but sharing a common topology and branch lengths. All trees were rooted using *Histoplasma mississippiense and H. capsulatum sensu stricto* (Sepúlveda et al. 2017).

#### Bayesian concordance analyses

In cases where speciation has occurred and genetic divergence has accrued, the phylogenetic signal across the whole genome should be congruent across loci. We measured the genome-wide genealogical concordance at a finer scale using a Bayesian concordance analysis (BCA). First, we identified orthologous genes using the BUSCO annotation pipeline (Simão et al. 2015; Waterhouse et al. 2018) which encompasses 1,316 Benchmarking Universal Single-Copy Orthologs. These group of genes have been curated in 25 different species of Ascomyceteous fungi (Pryszcz and Gabaldón, 2016; Waterhouse et al., 2018). Next, we used MrBayes v. 3.2.6 (Ronquist and Huelsenbeck 2003; Hhna et al. 2012). to generate posterior tree distributions for each single gene. We summarized the gene trees using the command *mbsum* in the BUCKy program (Ané 2010; Larget et al. 2010; Chung and Ané 2011). with a burn-in of 1,000 trees. We then fed individual gene genealogies to BUCKy v. 1.4.2, with four independent runs and four Markov chain Monte Carlo (MCMC) chains, each with 10 million generations with a burn-in period of 100,000. Five values of the α parameter (0.1, 0.5, 1, 5, 10) were tested, which correspond to the prior probability distribution for the number of distinct gene trees (Ané 2010). The level of support for each node is expressed as a concordance factor (CF), which ranges between 0 (no concordance between genealogies) and 1 (complete concordance). This approach allowed us infer the phylogenetic relationships between putative species, while estimating the genome-wide genealogical support for their monophyly.

### Genetic distance

To further assess the extent of genetic differentiation between phylogenetic species, we used the metric *π*_*INTER*_, the average number of nucleotide differences between one sequence randomly chosen from a population and another sequence randomly chosen from a second population or species. D_XY,_ or *π*_*INTER*_, followed the form:

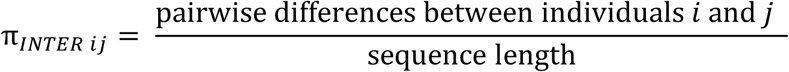

Mean *π*_*INTER*_ was the mean of all pairwise comparisons between individuals from two species. We calculated twenty mean *π*_*INTER*_values. We also calculated *π*_*INTER*_, the average pairwise genetic distance between individuals of the same species. *π*_*INTER*_ followed the same form as *π*_*INTER*_ but instead of calculating the average number of differences between species, it calculates the average number of differences between two randomly selected individuals of the same species. Mean *π*_*INTER*_ was the mean within-species value for each of the five species. We used Python for all calculations.

In cases were speciation is complete, *π*_*INTER*_ is expected to be much larger than *π*_*INTER*_. For each species pair, *π*_*INTER*_ can take two values (i.e., from each of the two species), so for the pairwise comparisons *π*_*INTER*_ is the pooled set of the two intraspecific distances. To compare the values of *π*_*INTER*_ and *π*_*INTER*_ for each species pair (10 pairwise comparisons), we used Two-Sample Fisher-Pitman permutation tests as implemented function *oneway_test* in the ‘*coin’* library in R (9,999 Monte Carlo resamplings; (Hothorn et al. 2008)). We also calculated *π*_*INTER*_ for each of the 10 pairwise comparisons and *π*_*INTER*_ for the five *Paracoccidioides* species for each of the largest 6 supercontigs.

### Gene exchange between species of *Paracoccidioides*

Previous work based on coding and microsatellite data suggested the possibility of gene exchange between *Paracoccidioides* species (Turissini et al. 2017). However, although microsatellite makers have the potential to reveal genealogical relationships between very closely related individuals, they are also prone to homoplasy as they mutate fast and their identity might not be caused by descent (Jarne and Lagoda 1996; Liepelt et al. 2001; Ellegren 2004). To address the possibility of gene exchange with better resolution, we used whole genome data and two complementary methods: *D*-statistics and *Int-HMM*.

### D-statistics

First, we calculated the excess of variants shared between potentially admixed species using D-statistics (Green et al. 2010; Durand et al. 2011; Martin et al. 2013, 2014). *D* is a metric to detect introgression from phylogenetic trees. The metric requires a four-taxon topology [(((P1,P2),P3),O)]. The allele in the outgroup (O) is labeled A, while the derived allele in the ingroup is labeled B. *D* compares the occurrence of two discordant site patterns, ABBA and BABA, representing sites in which an allele is derived in P3 relative to O, and is derived in one but not both of the sister lineages P1 and P2. These discordant patterns are most likely to arise if introgression occurs between P3 and either P2 or P1, in which case one site pattern will occur more frequently than the other. A positive *D* means introgression between P3 and P2; a negative *D* means introgression between P3 and P1. Due to the need a sorted topology, we focused on four species tetrads where [(((P1,P2),P3),O)]:

- [((*venezuelensis, restrepiensis*), *americana*), *lutzii*)]
- [((*venezuelensis, restrepiensis*), *brasiliensis*), *lutzii*)]
- [((*venezuelensis, brasiliensis*), *americana*), *lutzii*)]
- [((*restrepiensis, brasiliensis*), *americana*), *lutzii*)]

For each species pair, we measured the standard deviation of *D* from 1,000 bootstrap replicates. The observed genome-wide *D* was converted to a *Z*-score measuring the number of standard deviations it deviates from 0, and significance was assessed from a *P*-value using α = 0.01 as a cutoff after Holm–Bonferroni correction for multiple testing. We also calculated a variation of *D, f*_*D*_, which estimates the proportion of admixture by dividing the observed difference between the ABBA and BABA counts to the expected difference when the entire genome is introgressed. Besides the genome wide average of *D* and *f*_*D*_, we calculated both metrics for 5kb windows along the genome. We used DSuite for all calculations (Malinsky 2019), and used the allele frequencies within each species, as recommended in (Martin et al. 2014).

### Identification of introgressed haplotypes with Ancestry-HMM

we used a Hidden Markov Model (HMM), able to detect introgression in diploids and haploids (i.e., *Int-HMM;* (Turissini and Matute 2017a; Maxwell et al. 2018b,a)) to identify introgressed regions between all pairs of *Paracoccidioides* species. The HMM identifies introgressions between a pair of diverged populations or species: a donor, and a recipient (i.e., the admixed individual) by inferring the ancestry of every SNP in the genome. It then identifies a consecutive group of SNPs from the donor in the recipient background. Donor SNPs were selected such that they were monomorphic in the donor species and the allele frequency differences between the two species was greater than or equal to 30%. We also required that every individual in the donor species and at least one individual in the recipient species had a called genotype. Transition and emission probabilities of the HMM have been described elsewhere (Maxwell et al. 2018b,a).

### Identifying introgression tracts

*Int-HMM* determined the most probable genotype for each marker in each individual. We defined tracts as contiguous markers with the same genotype (species 1 or species 2). Introgressed SNPs are defined as those within a tract where the HMM probability for an introgression state (*d*) (i.e., originating from the donor) was ≥ 50%. In cases where we identified a region of *d* with at least 10 introgressed SNP markers flanked on one side by a small tract (under 10 SNPs) from the recipient that in turn was flanked by a single larger tract that was completely *d*, the two introgressed regions were merged and consolidated into a single tract. We did four consecutive rounds of filtering to allow identification of larger introgressed tracts that were broken up by small sections of the recipient species. These broken regions might be caused by gene conversion, double recombination events, or sequencing error (Figure S1 shows an example).

### Enrichment by sequence type

In cases where introgression is deleterious, selection will operate most efficiently against regions encoding functional elements (e.g. coding sequences, promoters; (Sankararaman et al. 2014; Juric et al. 2016; Turissini and Matute 2017a)). To test if a particular type of sequence was more or less prone to appear in introgressed regions, we partitioned the genome by sequence type into one of the following seven categories using the *P. brasiliensis* Pb18 genome annotations (ABKI00000000.2): CDS (coding sequence), 5’ -UTR, 3’ -UTR, intron, 2kb upstream inter (intergenic sequence 2kb upstream of a gene), 10kb inter (intergenic sequence within 10kb of a gene excluding 2kb upstream of a gene), and intergenic (intergenic sequence more than 10kb from a gene; (Desjardins et al. 2011)). Introgressions present in more than one individual but with different endpoints among isolates were broken into blocks, and these blocks were treated separately in the permutation test (described immediately below).

We calculated a summary statistic for each of the seven categories using the following definitions: ‘Introgressed percentage’ is the percentage of introgressions overlapping a given sequence type that occurred in any of the four possible introgression directions (two different species pairs and two reciprocal directions), ‘Genomic percentage’ is the proportion of the genome of any given type of sequence. ‘Enrichment’ is the ratio of the percentage of a given sequence type that has crossed species boundaries and the percentage of the genome encompassed by the same sequence type. We used permutation test in which each introgression block was randomly assigned to a new position in the genome to calculate *P-*values. For each permutation assay, we calculated the percentage of the randomly reassigned blocks overlapped with type of sequence. We repeated this procedure 10,000 times and generated a null-distribution for ‘Enrichment’. If introgressions are more likely to occur within a certain type of sequence than in the rest of the genome, ‘Enrichment’ will be greater than 1. Conversely, if introgressions are less likely to occur in a given type of sequence, ‘Enrichment’ will be less than 1. We determined whether introgressions were significantly enriched for any sequence type (i.e., a significant departure from 1), by comparing the observed ‘Enrichment’ and the distribution of resampled enrichments.

## RESULTS

### *Paracoccidioides* is haploid and shows little evidence for aneuploidy

We tested two aspects regarding the ploidy of *Paracoccidioides*. First, we assessed whether the coverage in the genome was homogenous. Significant deviations from homogeneity indicate aneuploidy. Second, we tested the global and local ploidy by scoring the mean coverage, and exploring for the presence of polymorphic sites. Haploid genomes, unlike higher ploidies, show no variation in each locus and thus there should show no minor—or major—alleles. Figure 1 shows the results of this analyses. First, the global distribution of coverage is distributed evenly around 1, with a few sites getting no coverage with respect to the reference genome (Figure 1A). Permutation tests show that this distribution is not significantly different from 1 (i.e., the expectation of homogenous ploidy along the genome; Two-Sample Fisher-Pitman Permutation Test, P = 0.078). When we ran similar analyses to study the mean coverage and minor allele frequency locally (i.e., in 5kb windows) along the *Paracoccidioides* genome. We found that the expectations of haploidy were fulfilled as well (i.e., no regions with abnormally high coverage, or that were systematically polymorphic; Figure 1B-E). Figures S2-S6 shows similar analyses for all species of *Paracoccidioides*. Overall these results confirm previous observations that *Paracoccidioides* is haploid (Almeida et al. 2007); for all analyses from now on, we treat *Paracoccidioides* as such.

**FIGURE 1.**
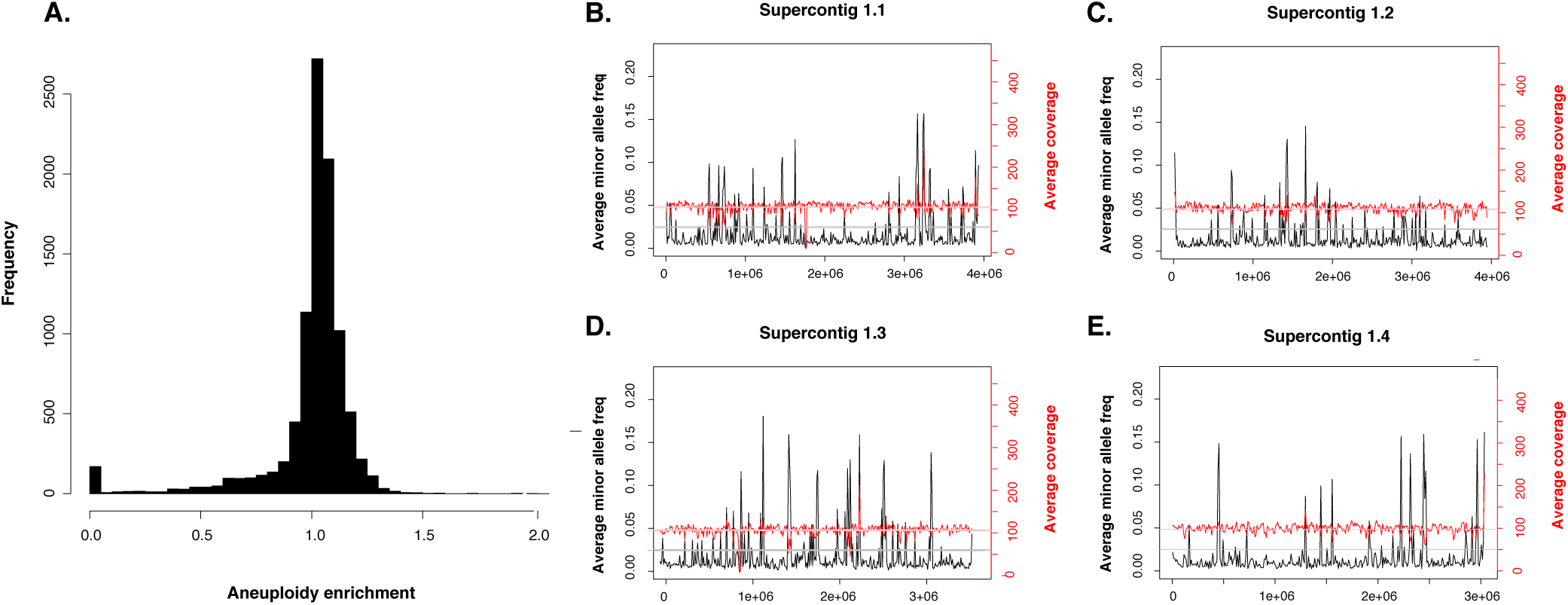
*Paracoccidioides* is haploid and shows no detectable levels of aneuploidy. **A.** Mean coverage per site shows no large and consistent deviations from the expected sequencing coverage suggesting little or no aneuploidy in *Paracoccidioides* genomes. **B.** Per site estimations of coverage (in red) show close adherence to the genome wide mean coverage for the whole genome. Per site estimations of the extent of polymorphism across the genome suggest that no single site shows evidence of polymorphism (cutoff: minor allele frequency > 20%) suggesting that besides no local aneuploidy, *Paracoccidioides* is haploid. The four panels on the right (B-E) show the coverage and minor allele frequency for the four largest supercontigs for the *P. lutzii* isolate EE.

### All proposed species show reciprocal monophyly and strong genealogical concordance

We evaluated the phylogenetic relationships between all species of the *Paracoccidioides* genus using genome wide variation and established how much of the genome supports these relationships. First, we built a maximum likelihood tree in which we used concatenated loci from the whole genome as the unit for phylogenetic analysis; (this approach has important limitations, (Mendes and Hahn 2018); see below). Figure 2A shows the resulting topology. Individual analyses of the 6 largest supercontigs yields similar topologies (Figure 2B-2G). Two results stand out from these analyses. As predicted by smaller efforts (Turissini et al. 2017), all five named species of *Paracoccidioides* are reciprocally monophyletic. The tree shows that all clusters previously described as species, including S1a and S1b, are highly supported by bootstrap and Bayesian probability independent on the analyzed scaffold. Similarly, genome-wide phylogenetic reconstruction shows similar results to previous approaches and suggest that *P. restrepiensis* and *P. venezuelensis* are the most closely related species of the group. *Paracoccidioides brasiliensis sensu stricto* is sister to the dyad *P. venezuelensis/P. restrepiensis*, and *P. americana* is an outgroup to the other three species of the *brasiliensis* species complex.

**FIGURE 2.**
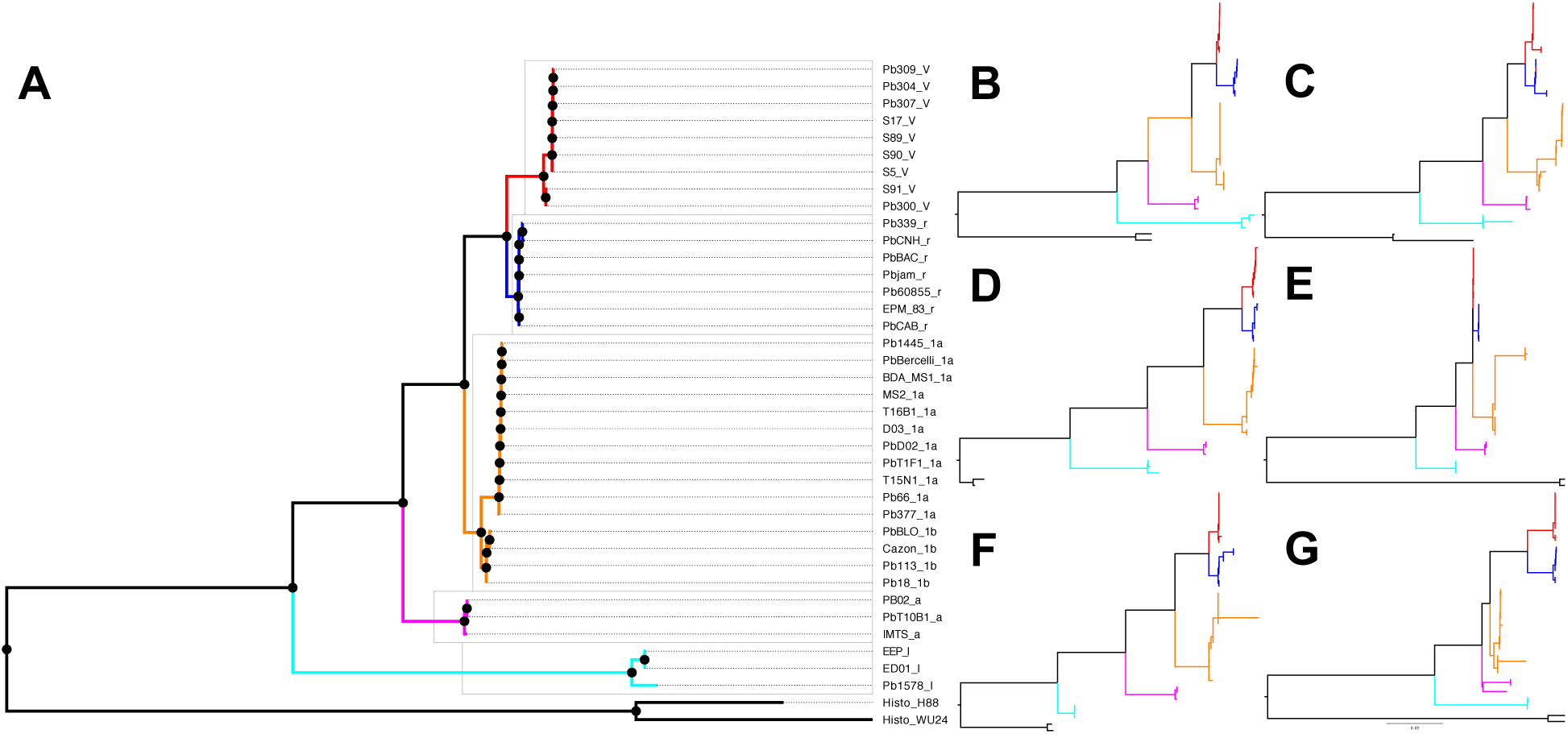
High level of differentiation between *Paracoccidioides* species. **A.** Maximum likelihood rooted phylogram using concatenated genome-wide loci. *Paracoccidioides venezuelensis* are marked in red and have a ‘V’ after the strain name. *Paracoccidioides restrepiensis* are marked in blue and have a ‘r’ after the strain name. *Paracoccidioides brasiliensis* are marked in orange and have a ‘1a’ or a ‘1b’ after the strain name. *Paracoccidioides americana* are marked in pink and have a ‘a’ after the strain name. *Paracoccidioides lutzii* are marked in cyan and have a ‘l’ after the strain name. **B-G**. Phylograms for the six largest supercontigs in the Pb18 genome show the same topology. We follow the same scheme as in Panel A. **B.** Supercontig 1.1. **C.** Supercontig 1.2. **D.** Supercontig 1.3. **E.** Supercontig 1.4. **F.** Supercontig 1.5. **G.** Supercontig 1.6. These analyses suggest that the species of *Paracoccidioides* are reciprocally monophyletic. The results are consistent to those shown in Muñoz et al. (2015, Figure 1) and Teixeira et al. (2020).

Additionally, we calculated the concordance factors for the each of the five *Paracoccidioides* species and the putative cryptic species S1a and S1b. If speciation has proceeded to extensive genetic differentiation, then most of the genome should show the signature of reciprocal monophyly in each of the proposed species. Figure 3 shows the results of a gene genealogy concordance analysis using BUCKy. The obtained topology is identical to that produced using maximum likelihood. The concordance factors for the five proposed species is in all cases over 90% which suggest that the vast majority of the genome shows concordance in the existence of the five species of *Paracoccidioides*. The only exceptions to this high level of concordance are the nodes that separates S1a and S1b which had a CF of 0.53 and 0.61. There is no metric on how high a CF must be to elevate a group to species level (Ané et al. 2006; Baum 2007), but the genome-wide concordance of this groups is much lower than the other *Paracoccidioides* species. For all analyses that follow, we treat *P. brasiliensis sensu stricto* as a single species with strong population structure.

**FIGURE 3.**
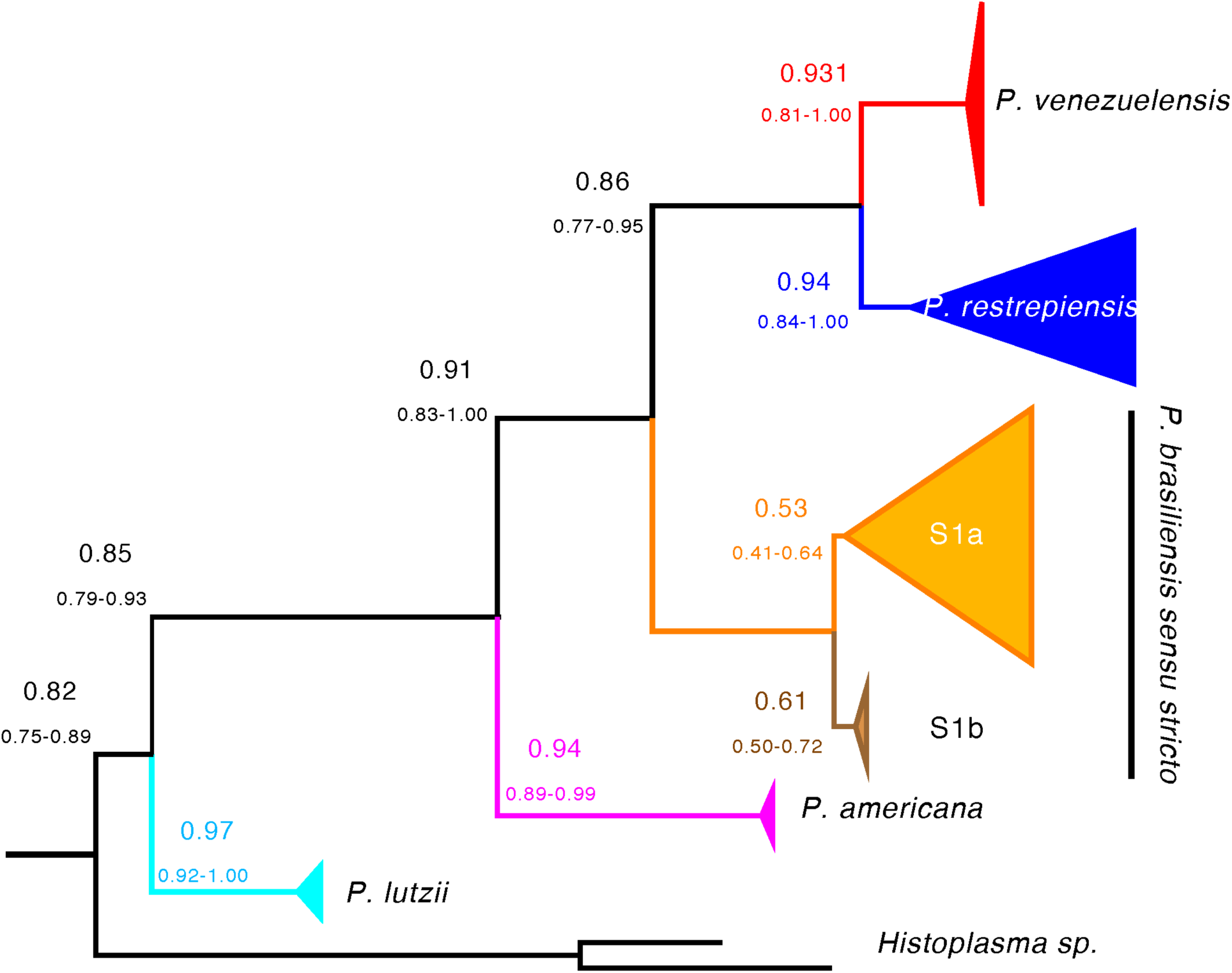
Species tree of the genus *Paracoccidioides* suggests that the majority of the genome of *Paracoccidioides* shows similar phylogenetic signal. The tree shown (built with BUCKy and a prior α = 5) is based on individual BUSCO genes. Values above each branch show the point estimate of the concordance factor (CF) for each node; values below branches show the 95% Bayesian credibility intervals for this estimate. The high levels of genomic concordance suggest reciprocal monophyly genome wide for the previously proposed species of *Paracoccidioides*.

Next, we calculated the mean genetic distance between individuals of the same species and between pairs of individuals from different species. The expectation is that pairwise comparisons between individuals from different species should show much higher differentiation than individuals from the same species (Hughes et al. 2009; Matute and Sepúlveda 2019). We found that genetic variability is partitioned among species and that in all cases the magnitude of interspecific distances is at least 2× higher than that of intraspecific distances for all species pairs in genome-wide estimations (Figure 4, Figures S7-S9). All pairwise comparisons between intraspecific and interspecific differentiation were significant (Two-Sample Fisher-Pitman Permutation Test, P < 1 × 10^−10^). As expected from the concordance analysis, the differentiation occurs along the whole genome (Figure 5, Figures S10-S12).

**FIGURE 4.**
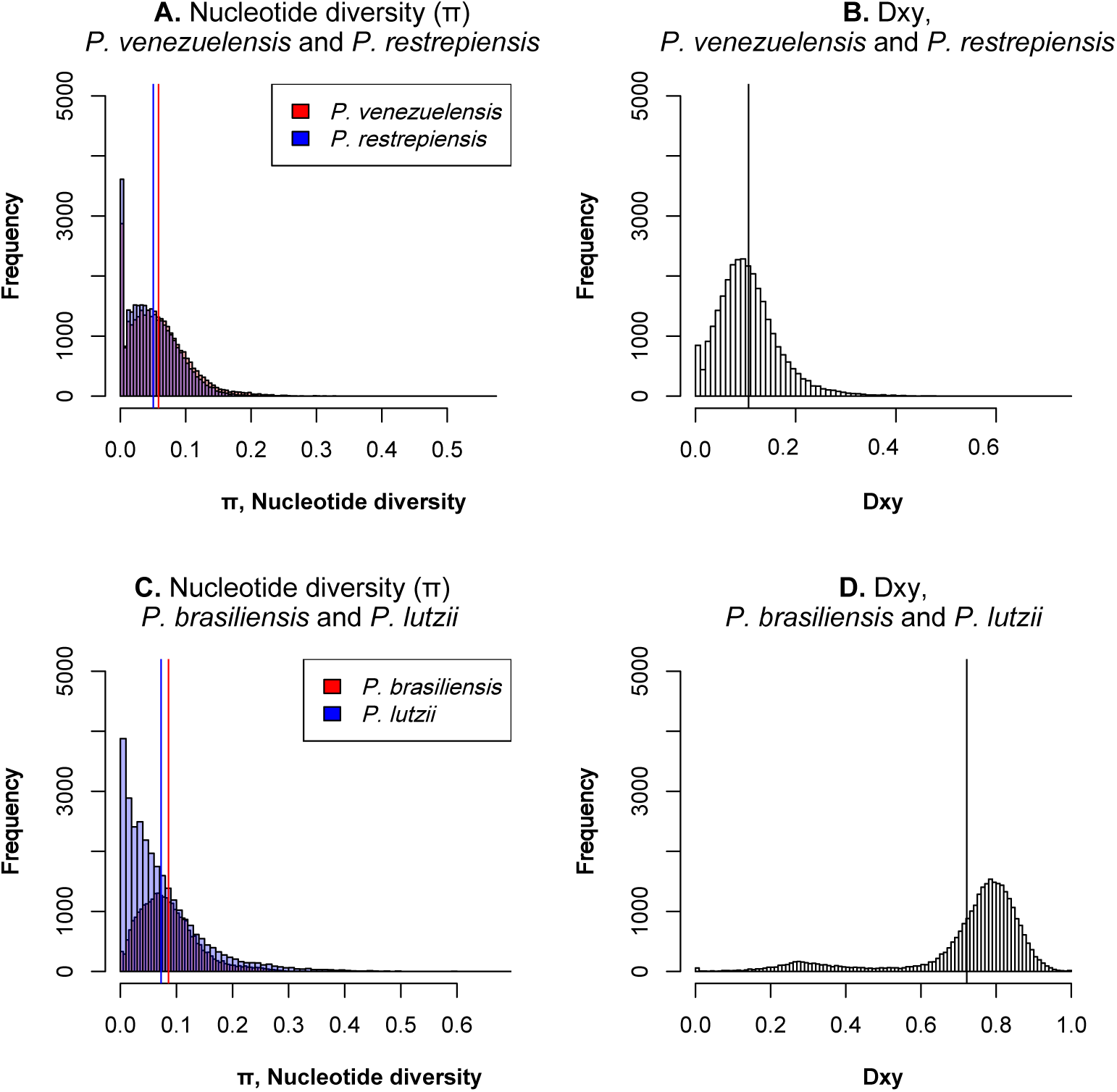
Genetic variation is partitioned across species in *Paracoccidioides*. Panels on the left (A, C) show the distribution of pairwise divergence within four species of *Paracoccidioides*. Blue and red bars show the two mean values of pi (one for each species). Panels on the left (B, D) show the distribution of average pairwise Dxy values between two species pairs: *P. venezuelensis* and *P. restrepiensis*, and *P. brasiliensis* and *P. lutzii*. Black lines show the mean value of pairwise Dxy. In all cases, the distributions of intraspecific and interspecific pairwise differences are non-overlapping (Two-Sample Fisher-Pitman Permutation Test, P < 1 × 10^−10^). Other species pairs are shown in Figure S7-S9.

**FIGURE 5.**
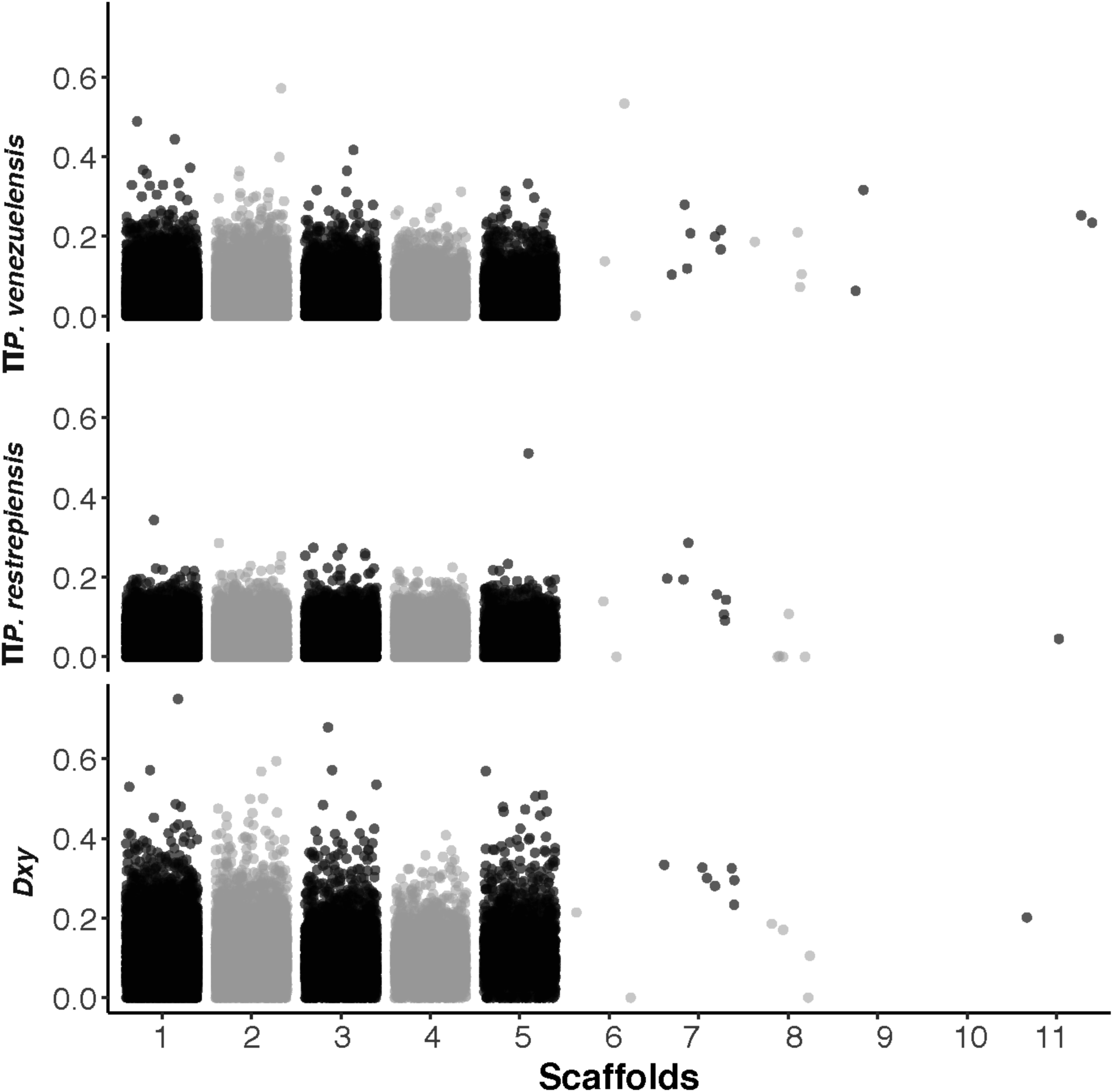
Differentiation between species of *Paracoccidioides* occurs genome-wide. π along the whole genome within *P. venezuelensis* (upper panel) and *P. restrepiensis* (middle panel) is lower than *Dxy* between species (upper panel). Other pairs of species are even more highly differentiated and are shown in Figures S7-S9.

The joint results from the phylogenetic analyses and the genetic distance calculations indicate that the genetic diversity within *Paracoccidioides* is partitioned across species. The five proposed species of *Paracoccidioides* fulfill the expectations of being advanced in the speciation continuum in terms of genomic divergence (Hendry et al. 2009; Roux et al. 2016). Next, we tested whether such divergence is accompanied by a reduction in the amount of gene flow between species.

### Low rates of detectable gene exchange between species

If speciation has proceeded to advanced stages, as seems to be the case for the species in *Paracoccidioides*, the magnitude of gene exchange between species should be limited. To detect potential alleles that have crosses species boundaries, we used two different methods: *D*-statistics and *Int-HMM*, a method that detects haplotypes likely to have crossed species boundaries. We describe the results for each species pair as follows.

#### *P. lutzii* and the species from the *brasiliensis* complex

Using *Int-HMM*, we found no evidence of introgression in any of these species pairs and in any reciprocal direction. If hybridization and admixture has occurred between *P. lutzii* and the other *Paracoccidioides* species, it has left no trace in the genomes of any of the involved species.

#### *P. americana* and the rest of species from the *brasiliensis* complex

*D* suggest that *P. americana* has donated more genetic material to *P. brasiliensis* and *P. venezuelensis* than it has donated to *P. restrepiensis*. In all cases the proportion of introgression is small (i.e., f_D_ is lower than 0.04, Table 1). We followed up with Ancestry-HMM and found no evidence of large haplotypes (over 500bp) between *P. americana* and the other species from the *brasiliensis* complex. This result suggest that introgressions are small and are probably old or strongly selected against. Regardless of the explanation for this low proportion of admixture, the joint results indicate that the magnitude of gene exchange between *P. americana* and the other species of the *brasiliensis* species complex is low.

**TABLE 1.**
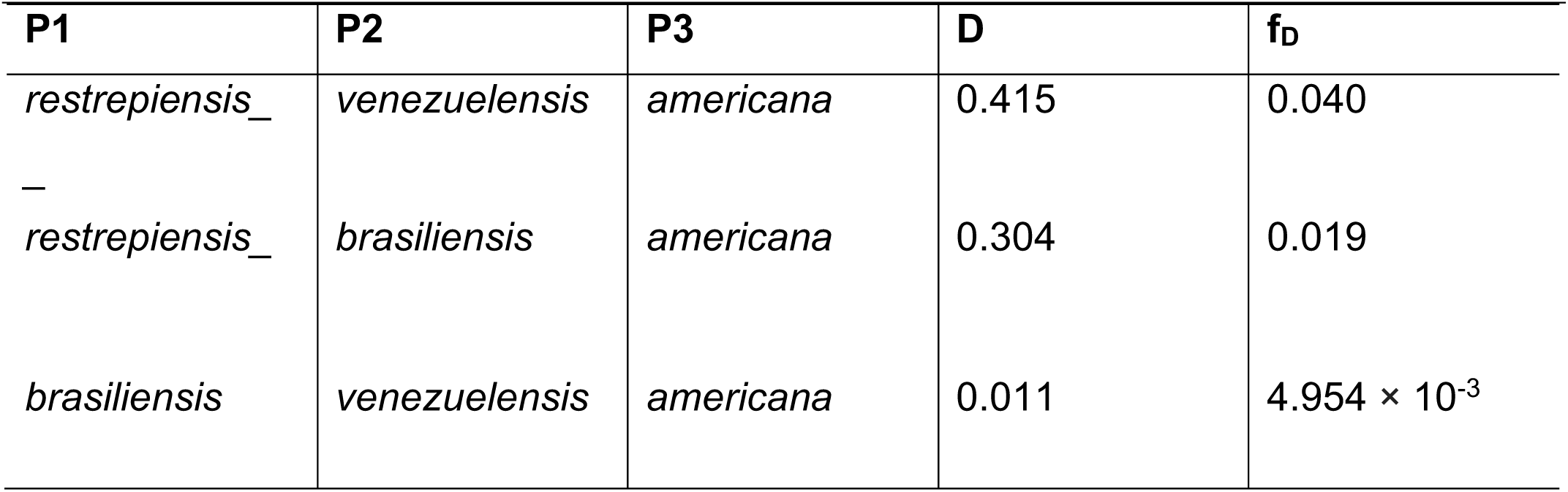
D and f_D_ values for species tetrads involving *P. americana* as a donor. In all cases, *P. lutzii* is the outgroup.

#### *P. brasiliensis/P. restrepiensis* and *P. brasiliensis/P. venezuelensis*

Unlike pairwise comparisons involving the more divergent species pairs, we found evidence of limited gene exchange using both methods in these two species pairs. The only tetrad that fulfilled the requirements for the calculation of *D* was [((*venezuelensis, restrepiensis*), *brasiliensis*), *lutzii*)]. The evidence of gene flow in this case was strong and showed that introgression between *P. brasiliensis* and *P. venezuelensis* is more common than *P. brasiliensis* and *P. restrepiensis* (D= 0.331, P <0.001; fd = 0.078). This amount of introgression is higher than those observed between *P. americana* and other species in the *brasiliensis* species group.

Next, we used *Int-HMM* to study the characteristics of the introgressed haplotypes. Table 2 shows the percentage of genome that has crossed species boundaries in each sequenced genome. We found no overlap of the introgressions regions between reciprocal directions or species pairs. In *P. brasiliensis/P. restrepiensis*, introgression mean length did not differ between reciprocal directions (Welch Two Sample t-test data: t = −0.857, df = 4.135, P = 0.438) and in both cases was ∼15kb, suggesting similar times of admixture or similar selection against introgression (Figure 6A). We found a similar pattern in *P. brasiliensis/P. venezuelensis.* The haplotype length did not differ between reciprocal directions (Welch Two Sample t-tests *brasiliensis-restrepiensis*: t = 0.449, df = 3.456, P = 0.680; *brasiliensis-venezuelensis*: t = 0.536, df = 50.795, P = 0.594) and in both cases was ∼15kb (Figure 6B). In all cases, introgressions are at low frequency (i.e., present in a single isolate per species) and mostly located in intergenic regions (Table 3). Figure 7 (A-D) shows the location of introgressed haplotypes in the two directions of the cross. Introgressions were distributed along most supercontigs suggesting that, most supercontigs are equally permissive—or refractory—to introgression (Figure 7).

**TABLE 2.**
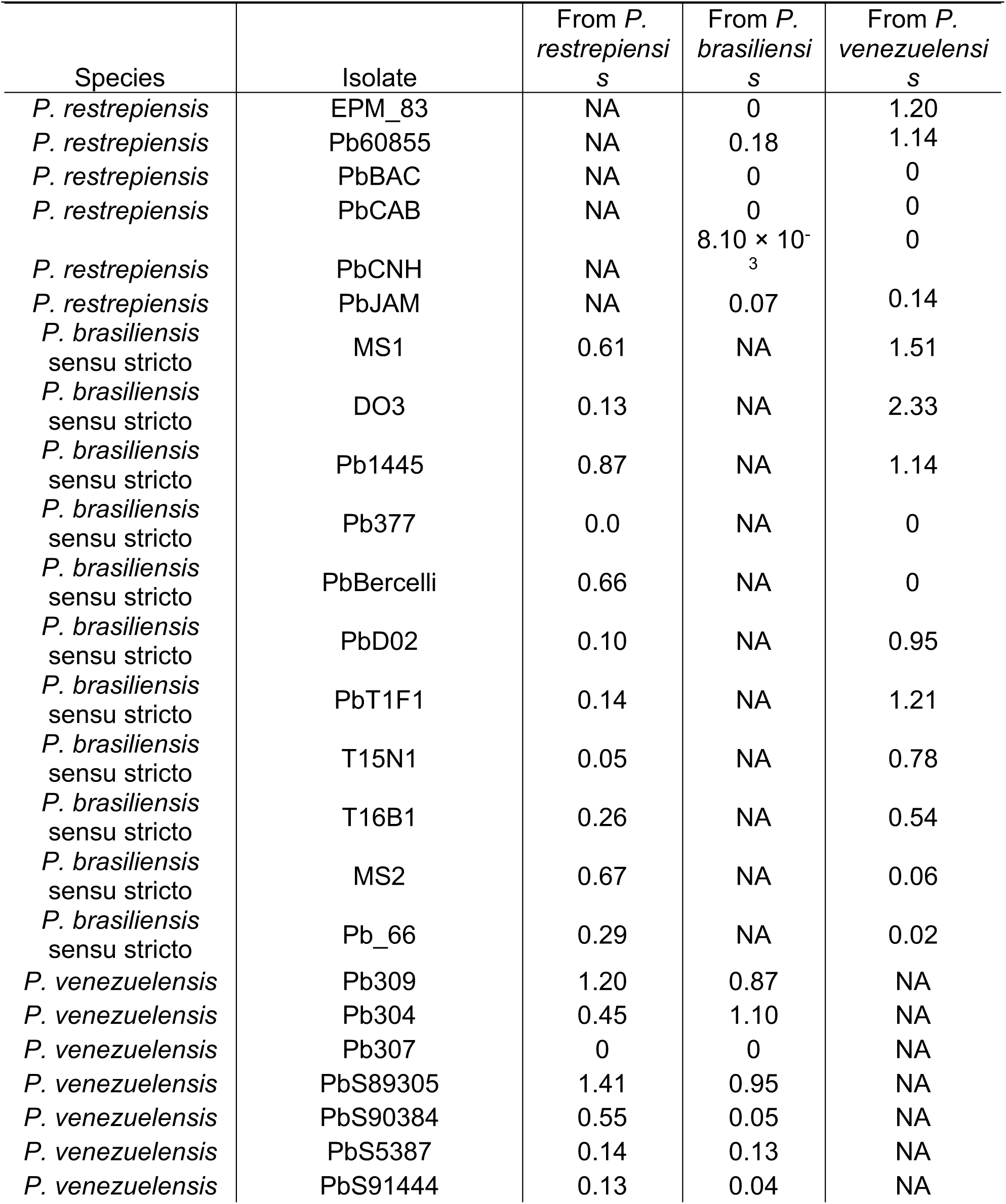
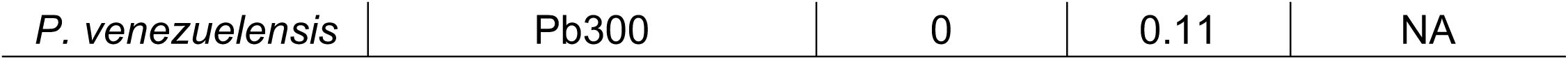
Percentage of the genome that shows introgression in each sequenced *P. restrepiensis, P. venezuelensis*, and *P. brasiliensis* isolate. Isolates from *P. lutzii* and *P. americana* show no evidence of large (over 500bp) introgressed haplotypes and are not listed.

**TABLE 3.**
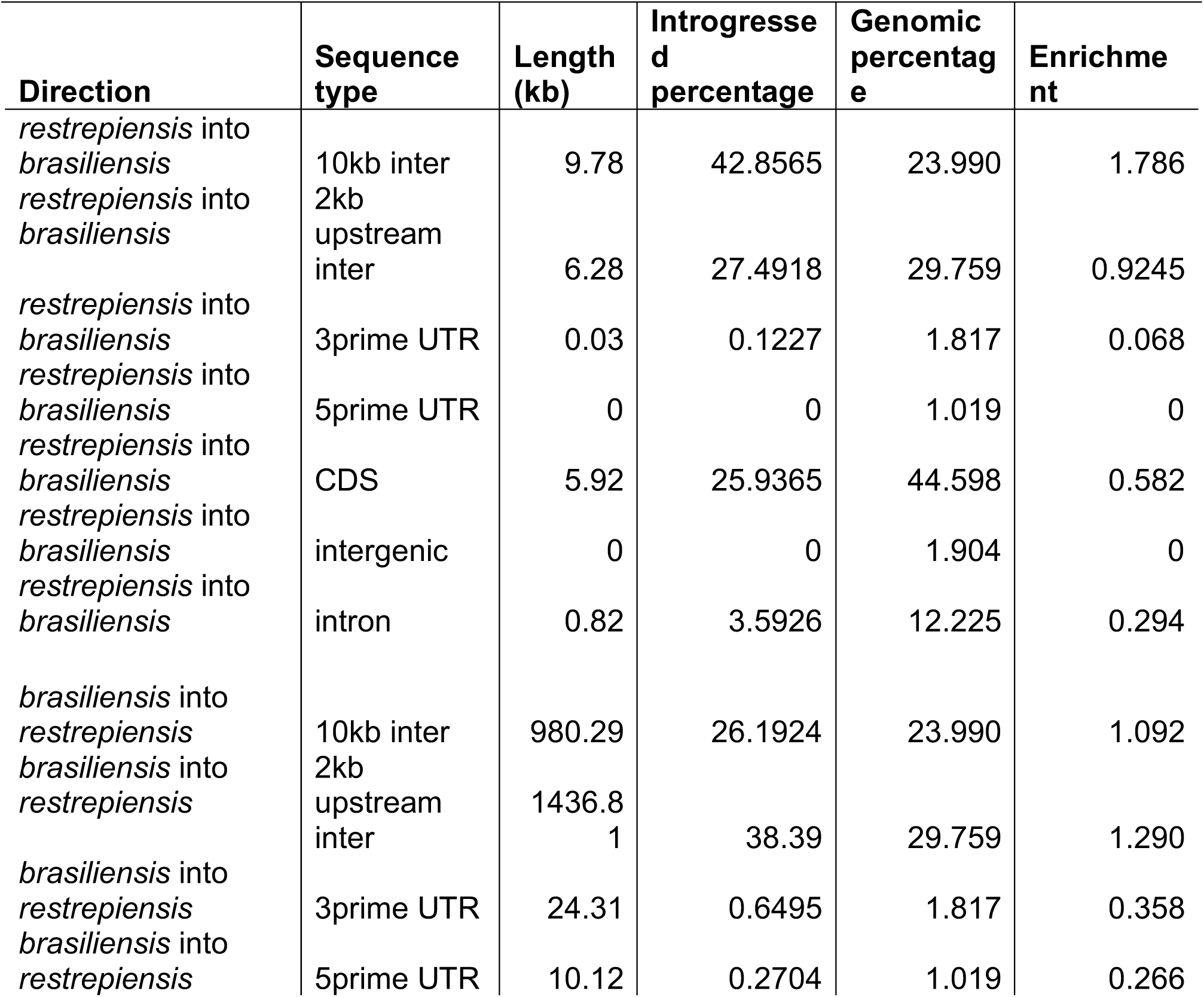
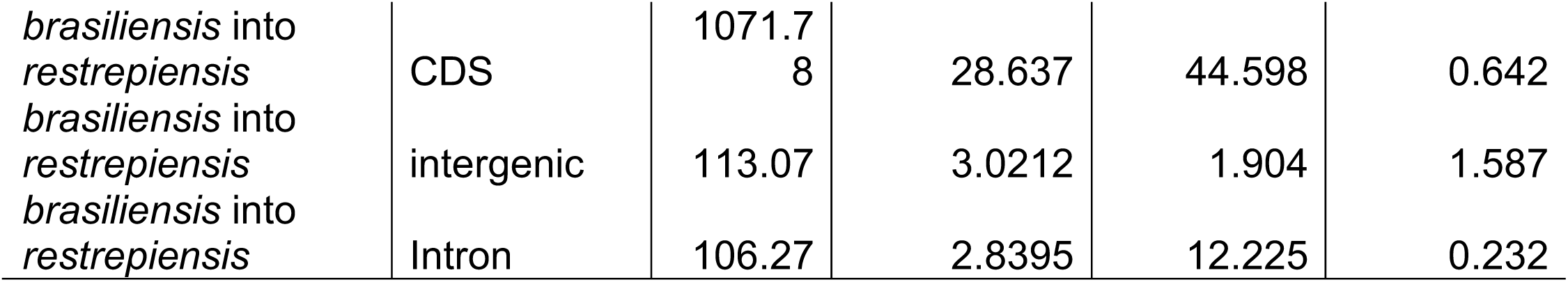
Introgressions between *P. restrepiensis* and *P. brasiliensis sensu stricto* are mostly found in intergenic regions. To study whether any particular type of sequence was over—or under—represented, we partitioned the genome by sequence type with each region being assigned to one of the following eight sequence types: CDS (coding sequence), exon, 5prime UTR, 3prime UTR, intron, 2kb upstream inter (intergenic sequence 2kb upstream of a gene), 10kb inter (intergenic sequence within 10kb of a gene), and intergenic (intergenic sequence more than 10kb from a gene). ‘Introgressed percentage’ is the percentage of introgressions overlapping a given sequence type for that direction, ‘Genomic percentage’ is the percentage of the genome represented by a given sequence type, and ‘Enrichment’ = (Introgressed percentage) / (Genomic percentage).

**FIGURE 6.**
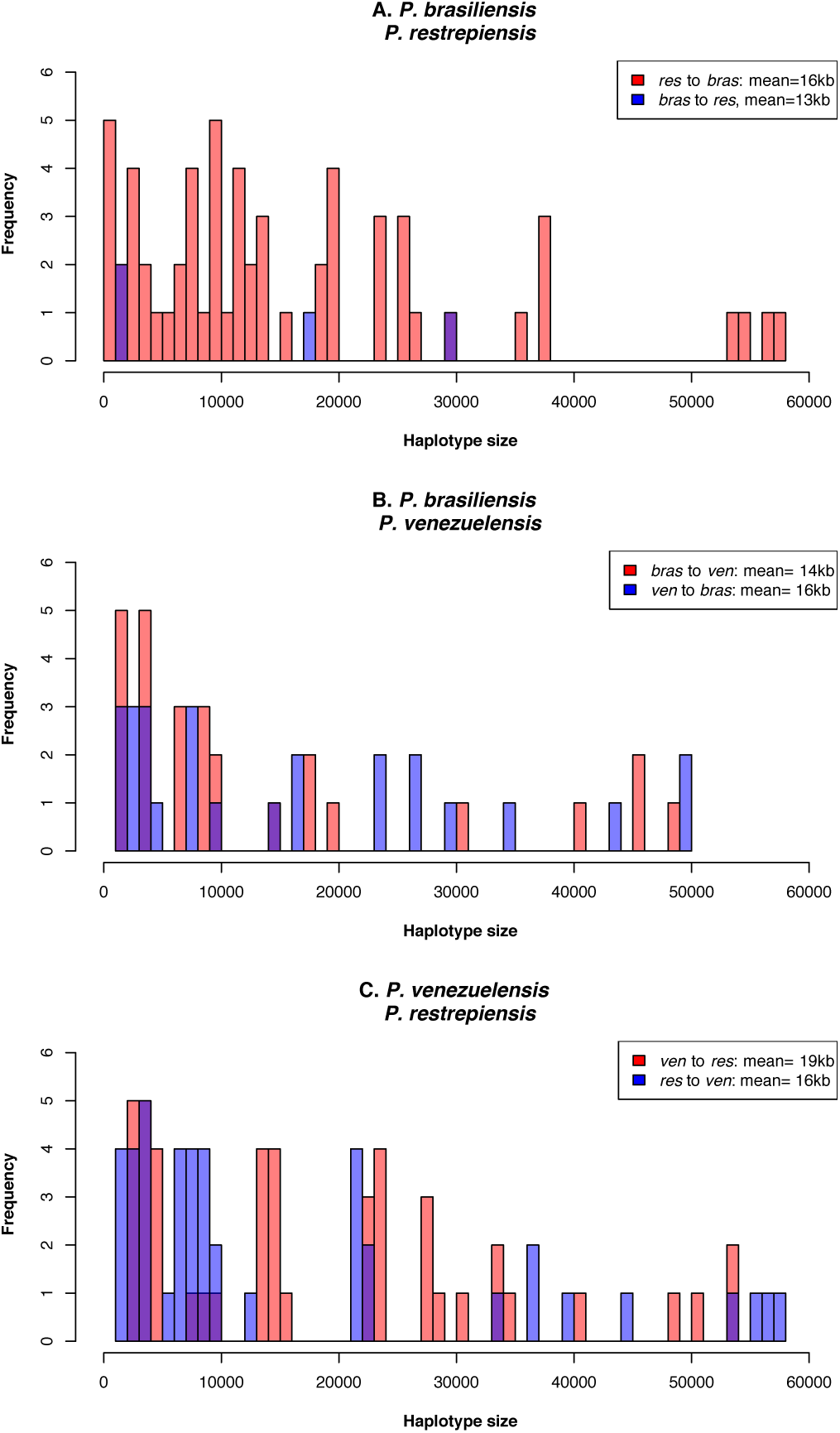
Distributions of the size of introgressed haplotypes in three species pairs of *Paracoccidioides* identified using *Int-HMM*. The *Int-HMM* algorithm was run on each individual separately and the distribution of haplotype sizes was computed for each of the two directions of introgression. **(A)** Size distribution of introgressed haplotypes between *P. brasiliensis sensu stricto* and *P. restrepiensis*. **(B)** Size distribution of introgressed haplotypes between *P. brasiliensis sensu stricto* and *P. venezuelensis*. **(C)** Size distribution of introgressed haplotypes between *P. venezuelensis* and *P. restrepiensis*. Even though there are large introgressions (over 50 kb) in most crosses, the majority of the introgressions are small.

**FIGURE 7.**
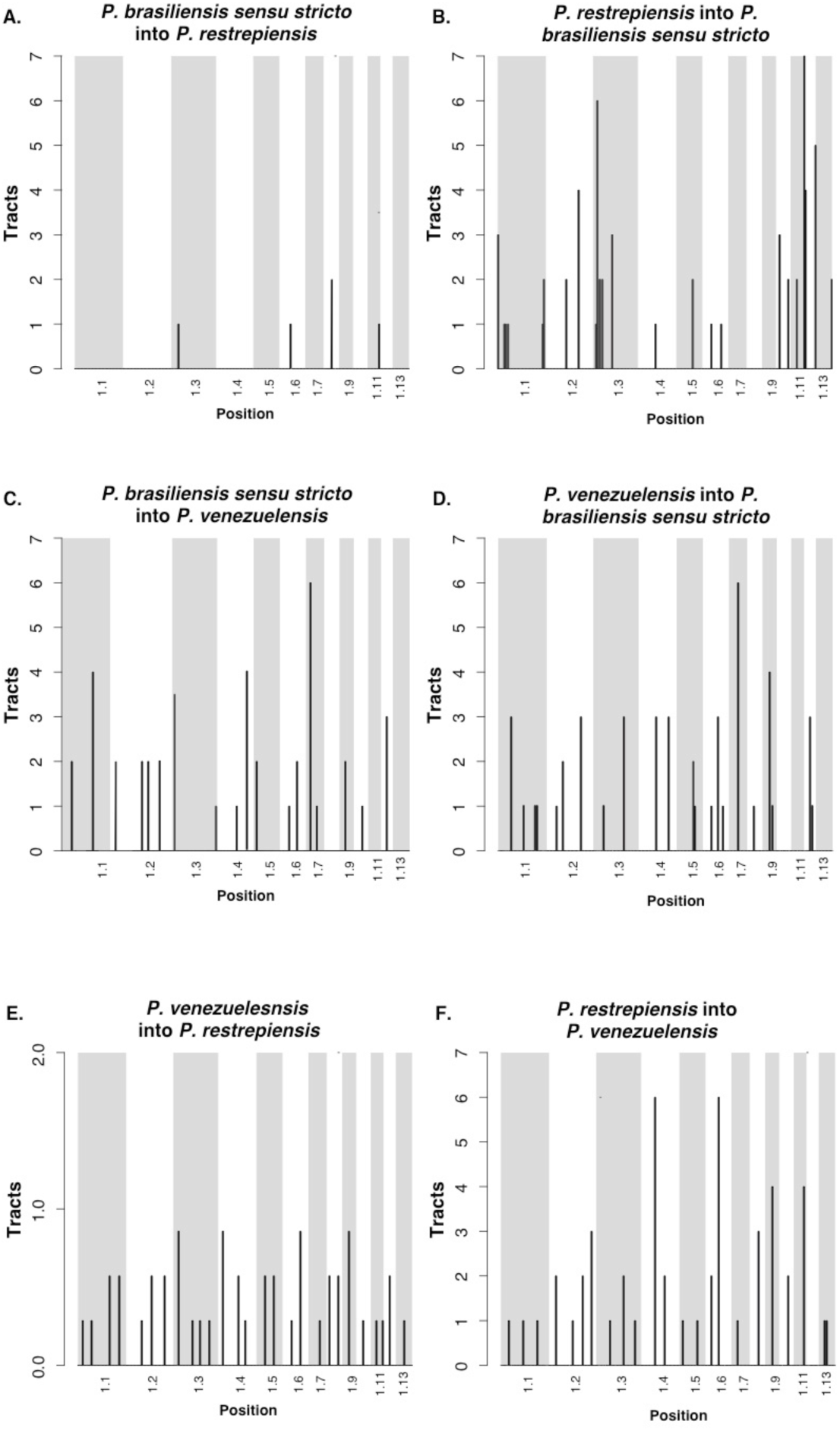
Location of the introgression tracts between species of *Paracoccidioides*. Each pair of panels shows the two reciprocal directions of introgression for a species pair. **A-B.** *P. restrepiensis* and *P. brasiliensis sensu stricto*. **C-D.** *P. venezuelensis* and *P. brasiliensis sensu stricto*. **E-F.** *P. restrepiensis* and *venezuelensis*. The two directions show differences in the amount of introgression, hence the differences in the scale of the *y*-axis. We plotted the largest 13 supercontigs of the genome. Histograms show the number of introgression tracts found 500 kb windows.

#### P. venezuelensis/P. restrepiensis

Finally, we studied the most recently diverged species pair in *Paracoccidioides* using *Int-HMM*. We found no overlap in the location of haplotypes between the two reciprocal directions or with any of the other dyads of *Paracoccidioides*. There was no difference in the genome proportion introgressed per individual between reciprocal directions (Welch Two Sample t-test t = 0.11, df = 9.95, P = 0.91). The mean haplotype length did not differ between reciprocal directions (Welch Two Sample t-test data: t = 0.741, df = 85.12, P = 0.461) and in both cases was ∼16kb (Figure 6C). As in the case for the other *Paracoccidioides* dyads, introgressions were at low frequency and largely in intergenic regions (Table 3). Notably, we found similar amounts of introgression in this pair as we found in the more divergent pairs (*venezuelensis*/*restrepiensis* vs. *venezuelensis*/*brasiliensis sensu stricto*: Welch Two Sample t-test, t = 1.205, df = 31.962, P = 0.237; *P. venezuelensis*/ *P. restrepiensis* vs. *P. restrepiensis*/*P. brasiliensis sensu stricto*: Welch Two Sample t-test, t = −1.372, df = 21.731, P = 0.184). Figure 6C shows the haplotype size frequency distribution of introgressions in the two reciprocal directions. Introgressions were distributed along the whole genome and did not follow a particular clustering pattern (Figure 7E, F).

## DISCUSSION

Our study uses genomic data to confirm previous observations that five species of *Paracoccidioides i*) are all haploid, and ii) are genetically differentiated. We also present results that suggest that the genomes of these species show strong levels of genealogical concordance genome wide and rarely exchange genes. The species of *Paracoccidioides* show considerable divergence and reciprocal monophyly which in turn suggest these five species are at an advanced stage on the speciation continuum (Hendry et al. 2009; Hudson and Coyne 2009; Roux et al. 2016).

Our analyses of the magnitude of gene flow between species confirm that even in spite of extensive geographic overlap, the *Paracoccidioides* species rarely exchange genes. In the more divergent pairs, those of the *brasiliensis* complex and *P. lutzii*, we found no evidence of introgression. This is consistent with the high levels of divergence between species which have been hypothesized to be over 30 million years apart (Teixeira et al. 2009, Turissini et al. 2017). We observed a similar—but not identical— pattern between *P. americana* and the other species of the *brasiliensis* complex (*brasiliensis, restrepiensis, venezuelensis*). Potential introgressions between these species are rare and of very small size making them potentially indistinguishable from incomplete lineage sorting as there was no evidence of gene exchange in any of the pairs. This paucity of gene exchange is not caused by lack of contact. *Paracoccidioides brasiliensis, P. lutzii*, and *P. americana* coexist in Brazil and have even been found in the same host (Matute et al. 2006; Teixeira et al. 2020). *Paracoccidioides venezuelensis* and *P. americana* share their geographic range in Venezuela as well. This extensive geographic overlap suggests that there is ample opportunity for gene exchange but it does not occur.

We do find evidence of moderate gene exchange in the triad *brasiliensis-restrepiensis-venezuelensis*. These low levels of gene exchange are consistent with advanced divergence among *Paracoccidioides* species. Our scans for gene exchange pose two additional questions. First, the rate of gene exchange is symmetrical in two *Paracoccidioides* species pairs: *P. brasiliensis/P. venezuelensis* and *P. venezuelensis/P restrepiensis*. The third species pair, *P. brasiliensis sensu stricto*/*P. restrepiensis*, shows strongly asymmetric introgression mostly found in intergenic regions. The reasons behind this pattern remain unknown. We formulate two possibilities. First, the direction of migration might be asymmetric between these two species. If *P. brasiliensis* migrants come into the range of *P. restrepiensis* more often than the reciprocal type of migration, then *P. brasiliensis* alleles should be found more frequently in the *P. restrepiensis* background that the reciprocal. A second possibility is that the *P. restrepiensis* background is less permissive of introgression because the introgressed alleles might have more deleterious effects. Since *P. restrepiensis* has a much smaller effective population size (Matute et al. 2006), then variants that might ameliorate the potentially deleterious effect of introgressed alleles should be more rare. The rates of migration, hybridization—or even intraspecific recombination— and of potential hybrid incompatibilities are unknown in *Paracoccidioides* and we cannot disentangle between these possibilities.

A second intriguing pattern is that *P. venezuelensis* and *P. restrepiensis* show a similar level of introgression as the other species pairs. As divergence increases, so should the number of incompatibilities (Matute et al. 2010; Moyle and Nakazato 2010; Wang et al. 2015) which in turn should reduce the proportion of genome that can flow from one species to the other (Muirhead and Presgraves 2016; Hamlin et al. 2020). Since the *venezuelensis*/*restrepiensis* dyad is more closely related than other species pairs within the *brasiliensis* complex, we expected a higher level of gene exchange. Our results do not support this hypothesis. Even though the precise reasons for this pattern remain unexplored, the role of geography might be of particular importance. The ranges of *P. venezuelensis* and *P. restrepiensis* are contiguous but have not been reported to overlap. This differs from all other species pairs in *Paracoccidioides* which show some degree of geographic overlap (Teixeira et al. 2020). A precise assessment of the range and opportunity for hybridization will be crucial to establish the genetic, environmental, and demographic factors that govern the patterns of introgression in *Paracoccidioides*.

The identification of species boundaries and introgression in fungal pathogens has human health-related implications. *Paracoccidioides lutzii* and *P. brasiliensis sensu stricto* show differences in the immunological response they elicit (Lenhard-Vidal et al. 2013; Siqueira et al. 2016), the strength of the disease they cause (Siqueira et al. 2016), and in traits involved in diagnostic tools (De Almeida et al. 2015; Arantes et al. 2017; Comparato Filho et al. 2019). Introgression, then, can be a vehicle to transfer virulence factors and antifungal resistance in *Paracoccidioides*. Gene exchange can also be an source of variation in other fungal pathogens (Desjardins et al. 2017; Maxwell et al. 2018a,b). A systematic survey to characterize the virulence and resistance of differentiated species could reveal the extent to which diversification of the ethological agents of PCM has also led to divergence in virulence strategies. The combination of phenotypic studies and population genetics can also reveal whether gene exchange plays a role on the transfer of virulence factors and antifungal resistance strategies. Our results are in line with other studies that show gene exchange in fungi but also reveal that introgression is not an unavoidable outcome of secondary contact.

Geographic overlap is not synonymous with hybridization and in cases of diverged species (such as *P. lutzii* and the species of the *brasiliensis* species complex), hybrids might not occur even when species share a close geographic range. Hybrids might also be sterile or inviable (Giraud and Gourbière 2012). Genome factors such as the amount of divergence between hybridizing species (Hamlin et al. 2020), the landscape of recombination (Schumer et al. 2018; Martin et al. 2019) affect whether an introgression persists after hybridization. The different levels of divergence and the ample opportunity for hybridization among *Paracoccidioides* species provide for a system to test the relative importance of genomic factors on determining the amount of introgression occurs in nature.

## ACKNOWLEDGEMENTS

We would like to thank the members of the Matute lab for helpful scientific discussions and comments. M.M.T was supported by Conselho Nacional de Ciência e Tecnologia (CNPq) under contract 43460/2018-2.. This work was supported by NIH award R01GM121750. The authors have no conflicts of interest.

## SUPPLEMENTARY MATERIAL

**TABLE S1.**
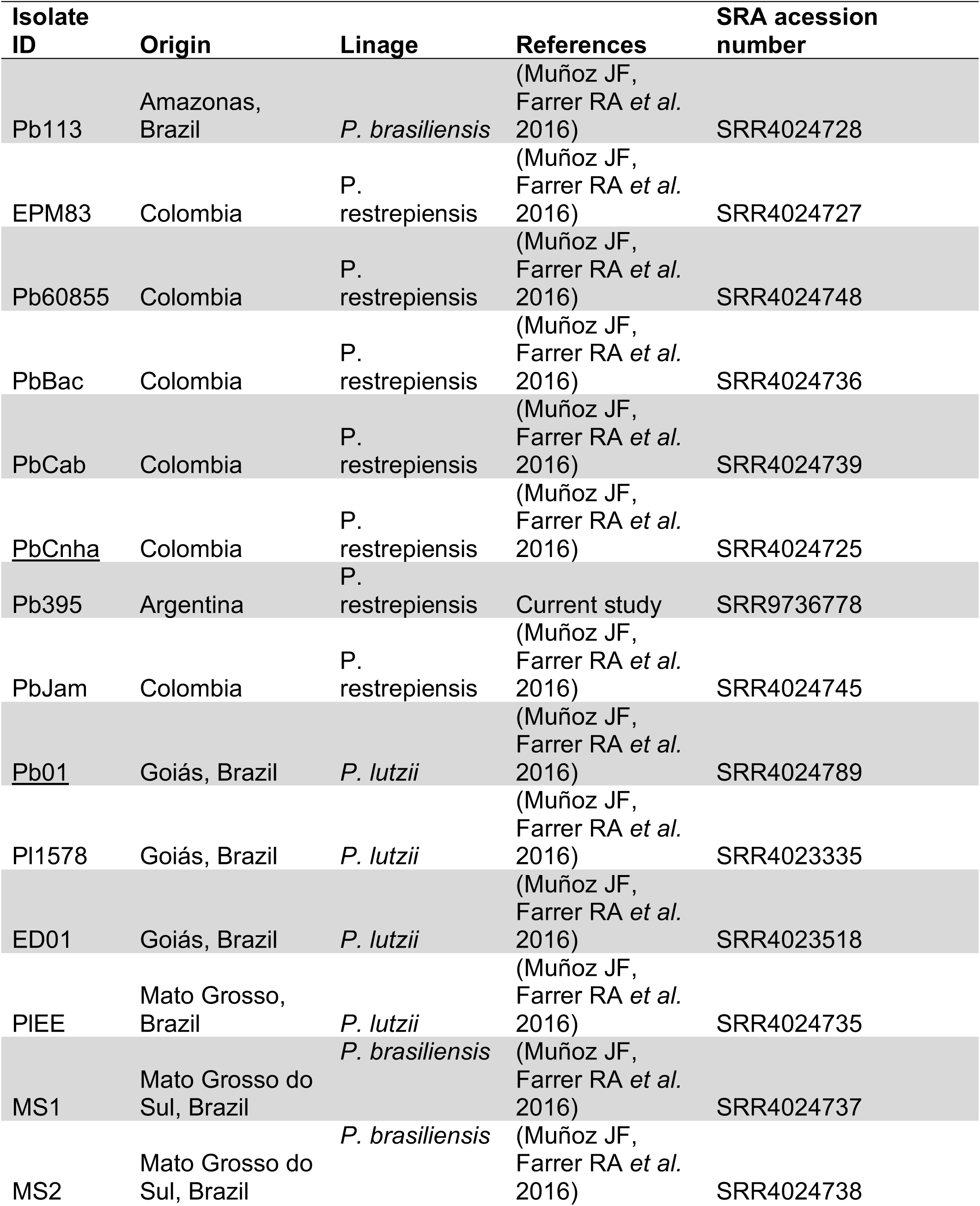
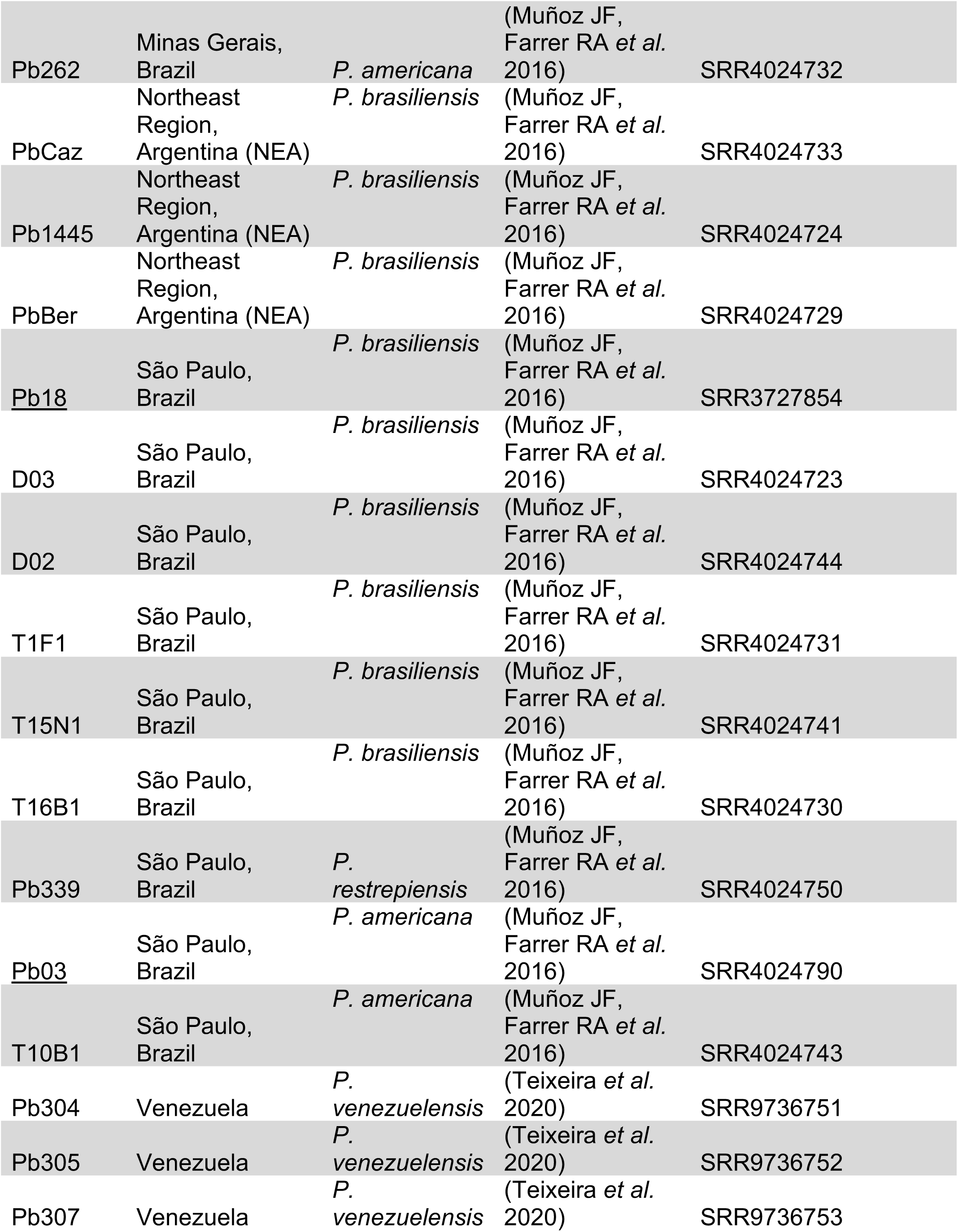
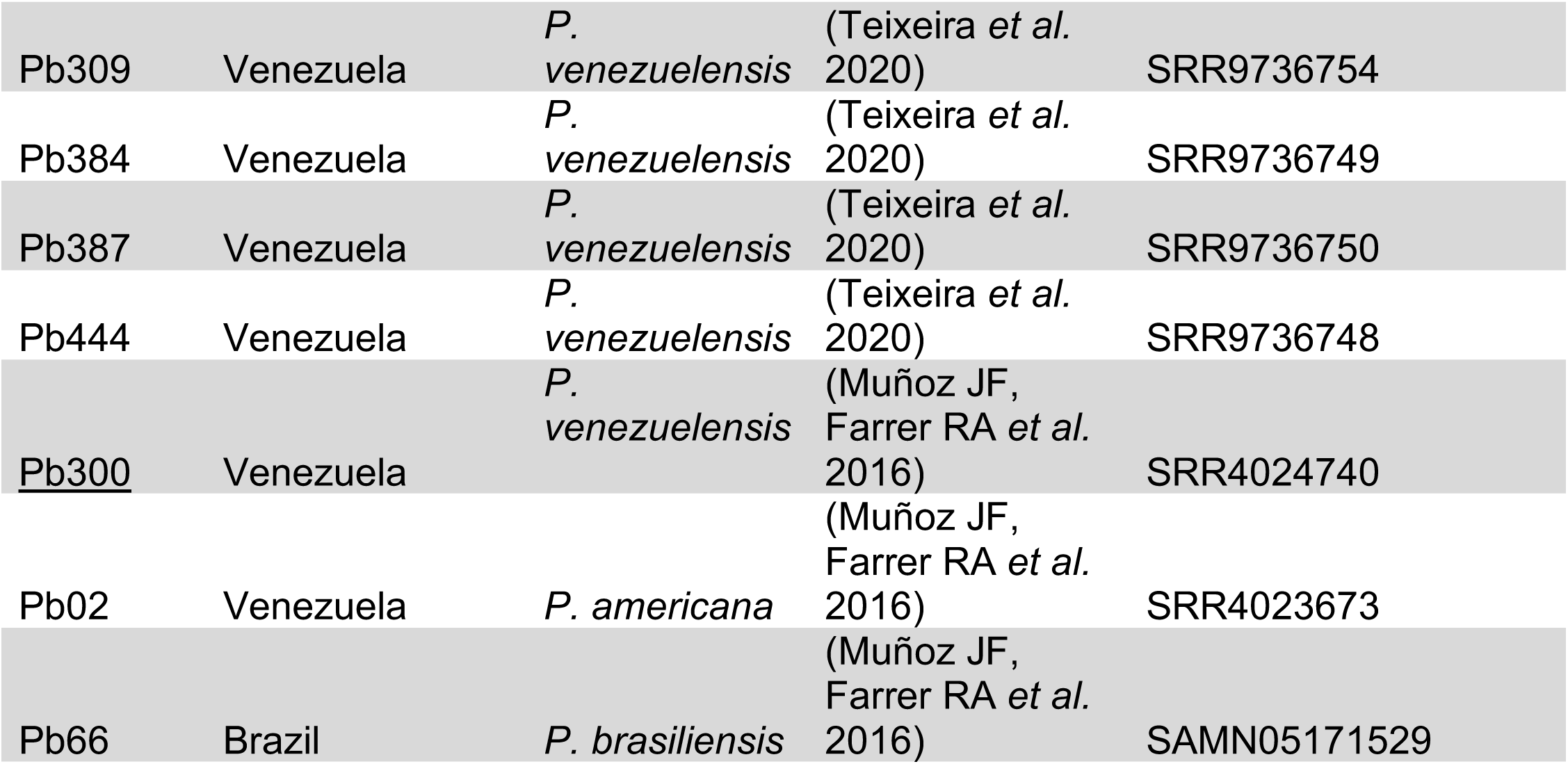
SRA numbers of the genomes used in this manuscript.

**FIGURE S1.**
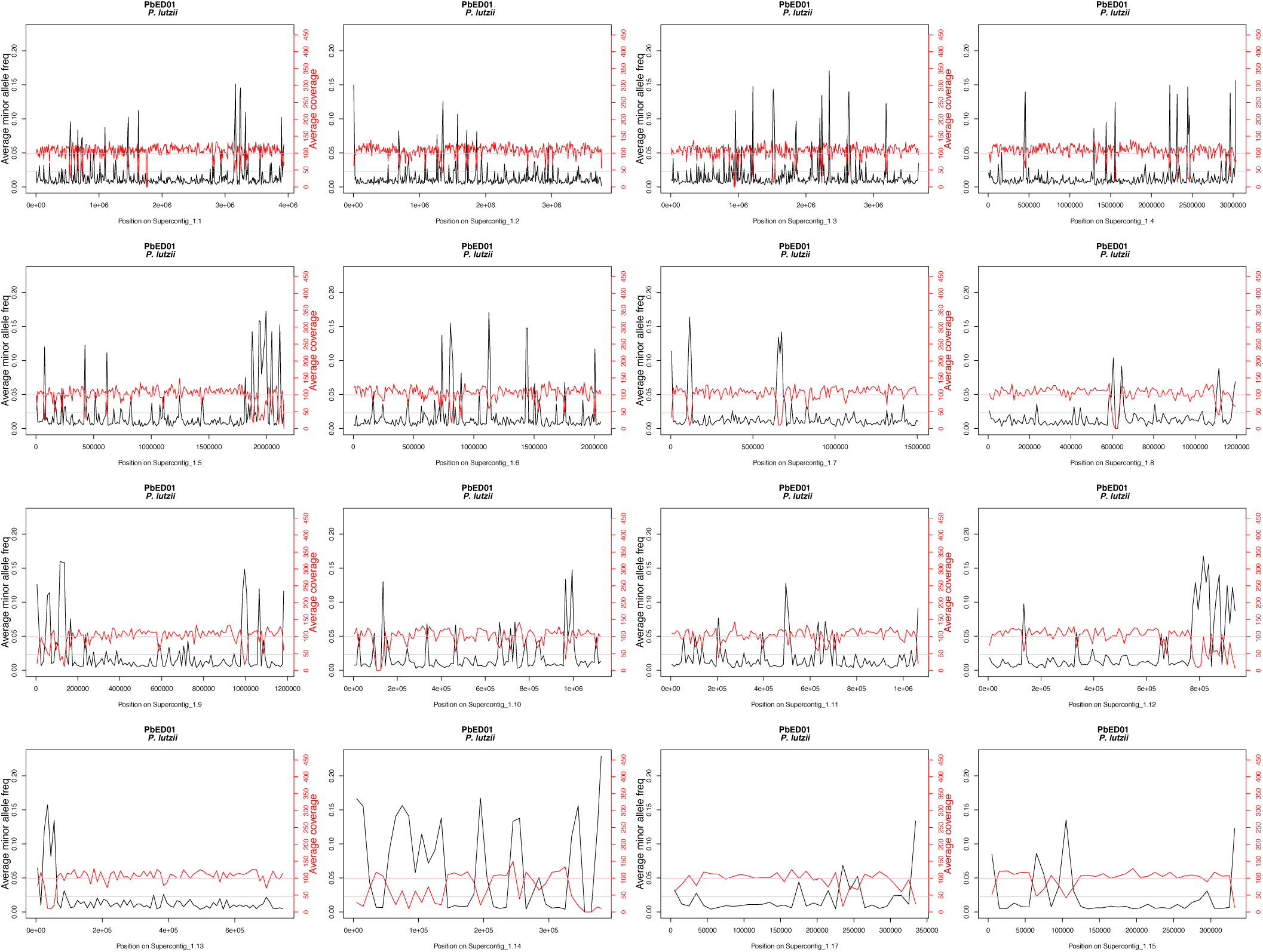
*Paracoccidioides lutzii* is haploid and shows no detectable levels of aneuploidy. Coverage and minor allele frequency plots for the largest 16 supercontigs. Per site estimations of coverage (in red) show close adherence to the genome wide mean coverage for the whole genome. Per site estimations of the extent of polymorphism across the genome suggest that no single site shows evidence of polymorphism (cutoff: minor allele frequency > 20%) suggesting that besides no local aneuploidy, *Paracoccidioides* is haploid. The plot shows results for isolate ED01; plots for other lines can be found in the Dryad package (doi: TBD).

**FIGURE S2.**
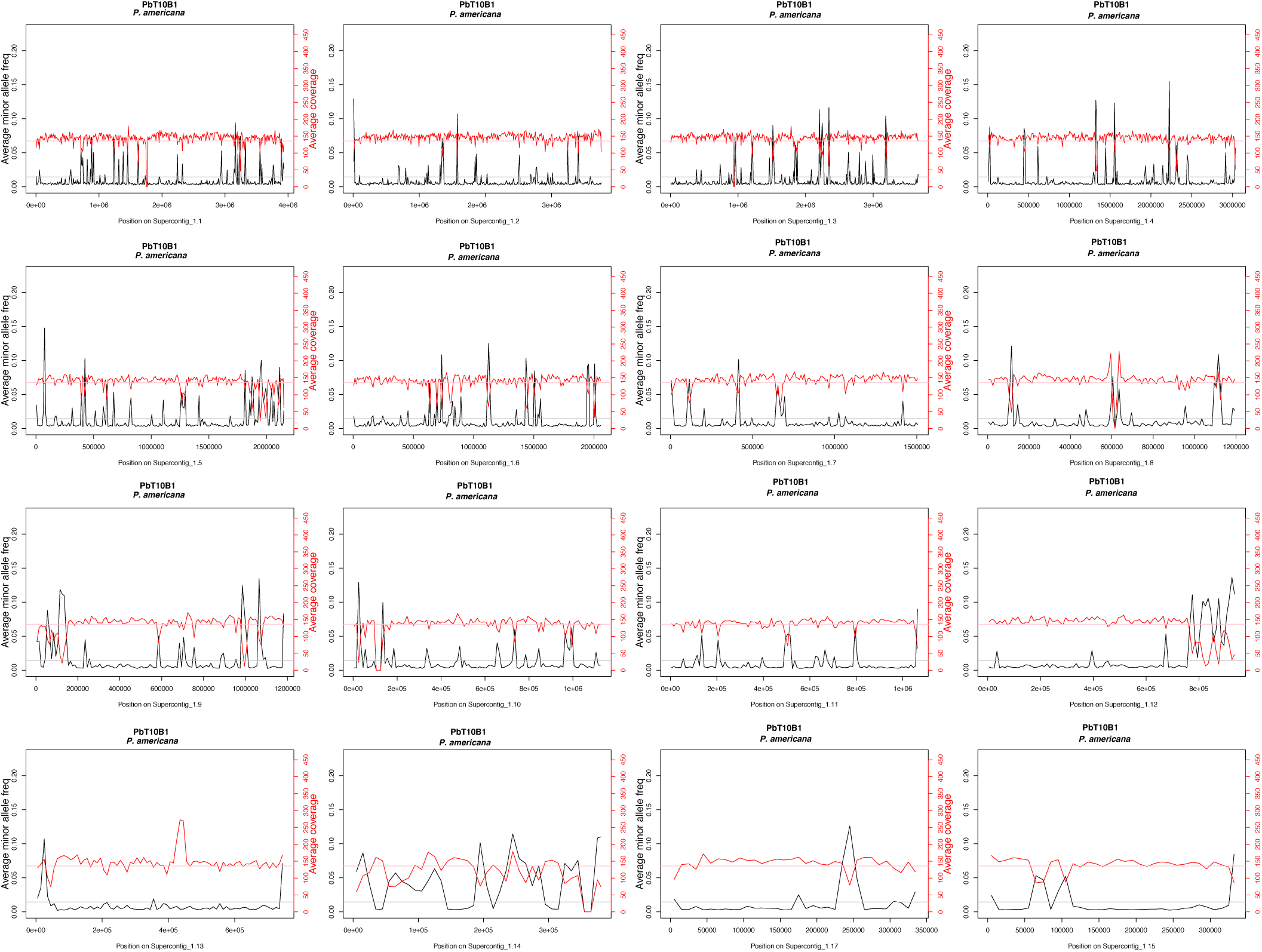
*Paracoccidioides americana* is haploid and shows no detectable levels of aneuploidy. Coverage and minor allele frequency plots for the largest 16 supercontigs. Per site estimations of coverage (in red) show close adherence to the genome wide mean coverage for the whole genome. Per site estimations of the extent of polymorphism across the genome suggest that no single site shows evidence of polymorphism (cutoff: minor allele frequency > 20%) suggesting that besides no local aneuploidy, *Paracoccidioides* is haploid. The plot shows results for isolate T10B1; plots for other lines can be found in the Dryad package (doi: TBD).

**FIGURE S3.**
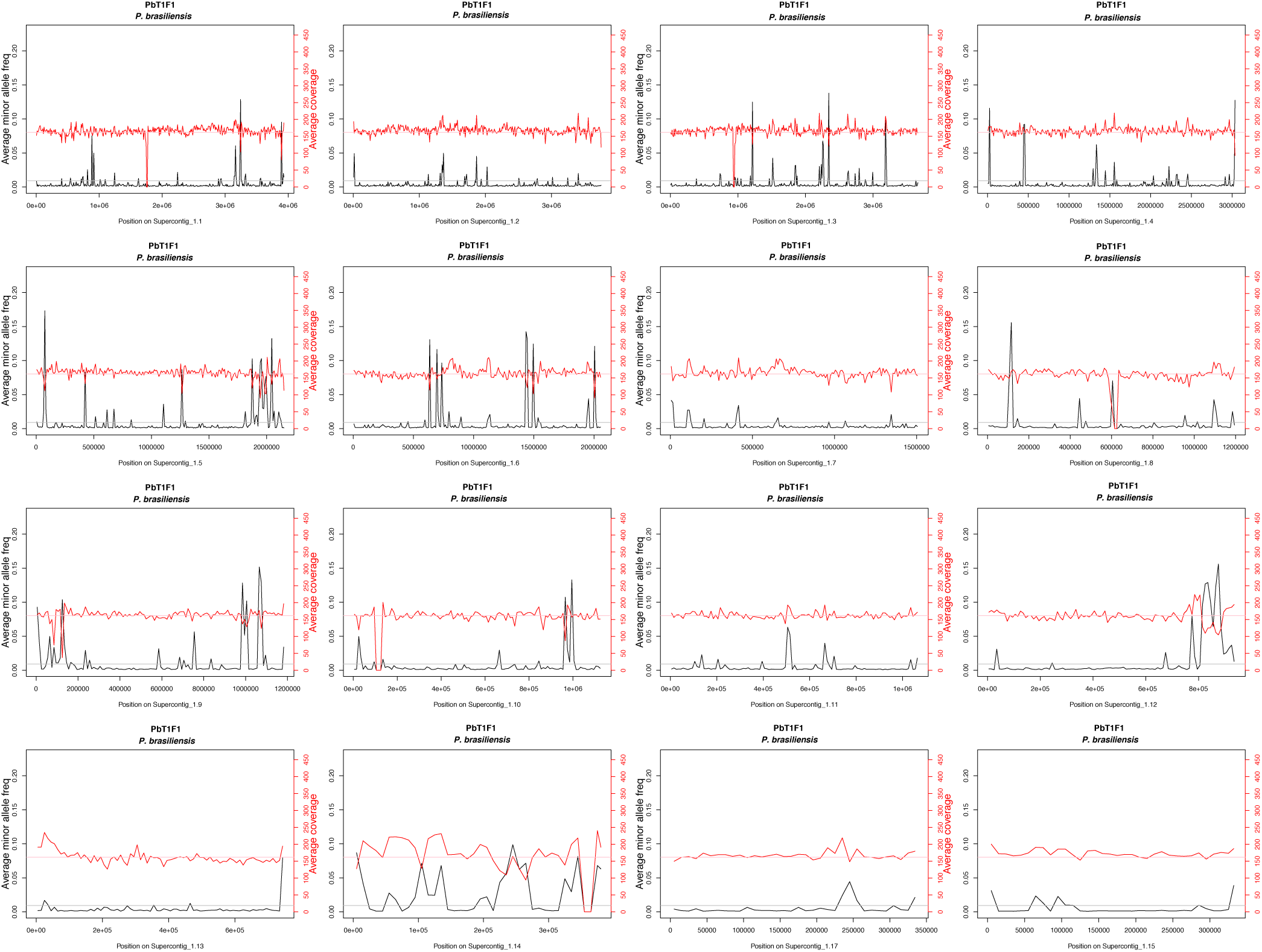
*Paracoccidioides brasiliensis* is haploid and shows no detectable levels of aneuploidy. Coverage and minor allele frequency plots for the largest 16 supercontigs. Per site estimations of coverage (in red) show close adherence to the genome wide mean coverage for the whole genome. Per site estimations of the extent of polymorphism across the genome suggest that no single site shows evidence of polymorphism (cutoff: minor allele frequency > 20%) suggesting that besides no local aneuploidy, *Paracoccidioides* is haploid. The plot shows results for isolate T1F1; plots for other lines can be found in the Dryad package (doi: TBD).

**FIGURE S4.**
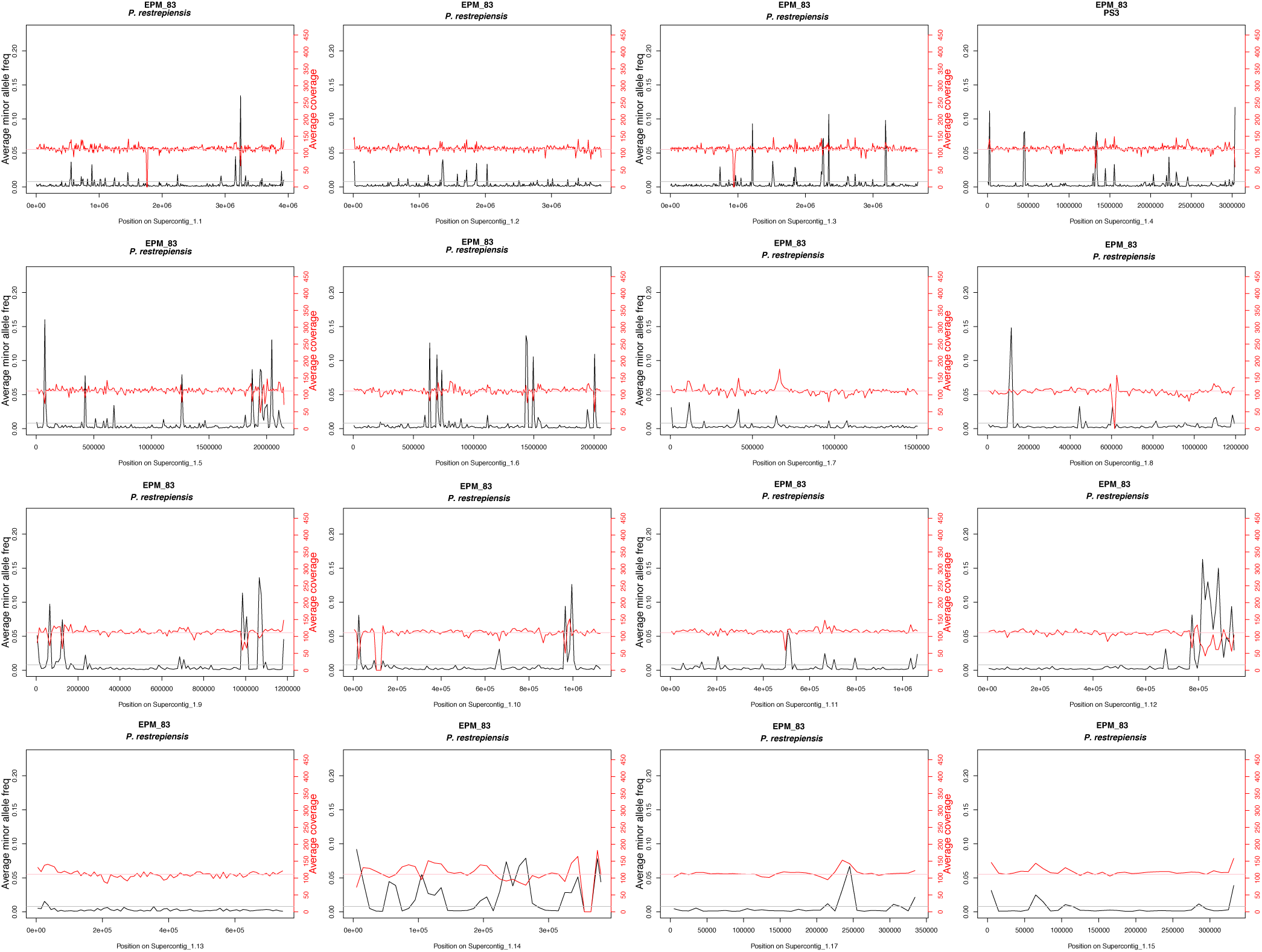
*Paracoccidioides restrepiensis* is haploid and shows no detectable levels of aneuploidy. Coverage and minor allele frequency plots for the largest 16 supercontigs. Per site estimations of coverage (in red) show close adherence to the genome wide mean coverage for the whole genome. Per site estimations of the extent of polymorphism across the genome suggest that no single site shows evidence of polymorphism (cutoff: minor allele frequency > 20%) suggesting that besides no local aneuploidy, *Paracoccidioides* is haploid. The plot shows results for isolate EPM83; plots for other lines can be found in the Dryad package (doi: TBD).

**FIGURE S5.**
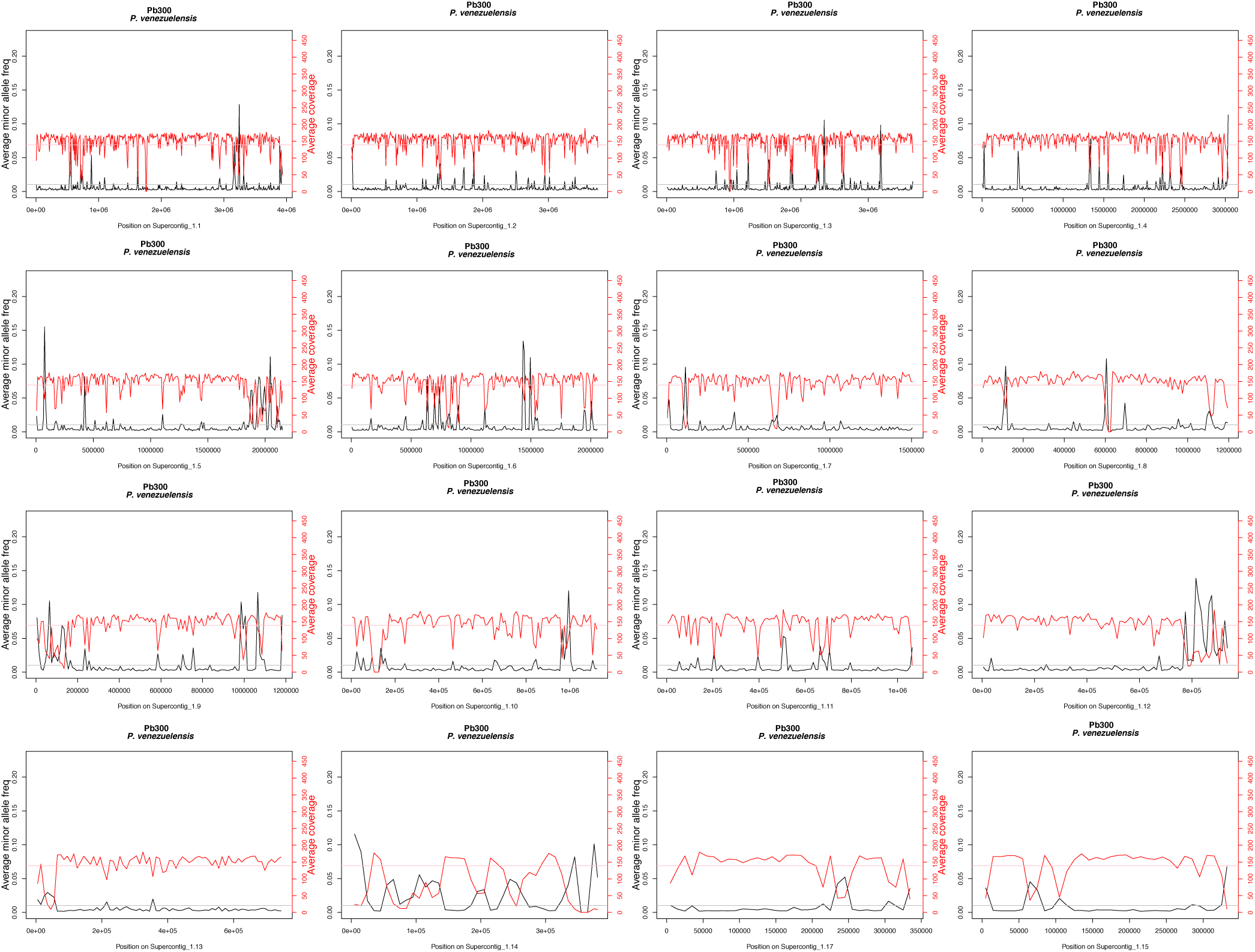
*Paracoccidioides venezuelensis* is haploid and shows no detectable levels of aneuploidy. Coverage and minor allele frequency plots for the largest 16 supercontigs. Per site estimations of coverage (in red) show close adherence to the genome wide mean coverage for the whole genome. Per site estimations of the extent of polymorphism across the genome suggest that no single site shows evidence of polymorphism (cutoff: minor allele frequency > 20%) suggesting that besides no local aneuploidy, *Paracoccidioides* is haploid. The plot shows results for isolate Pb300; plots for other lines can be found in the Dryad package (doi: TBD).

**FIGURE S6.**
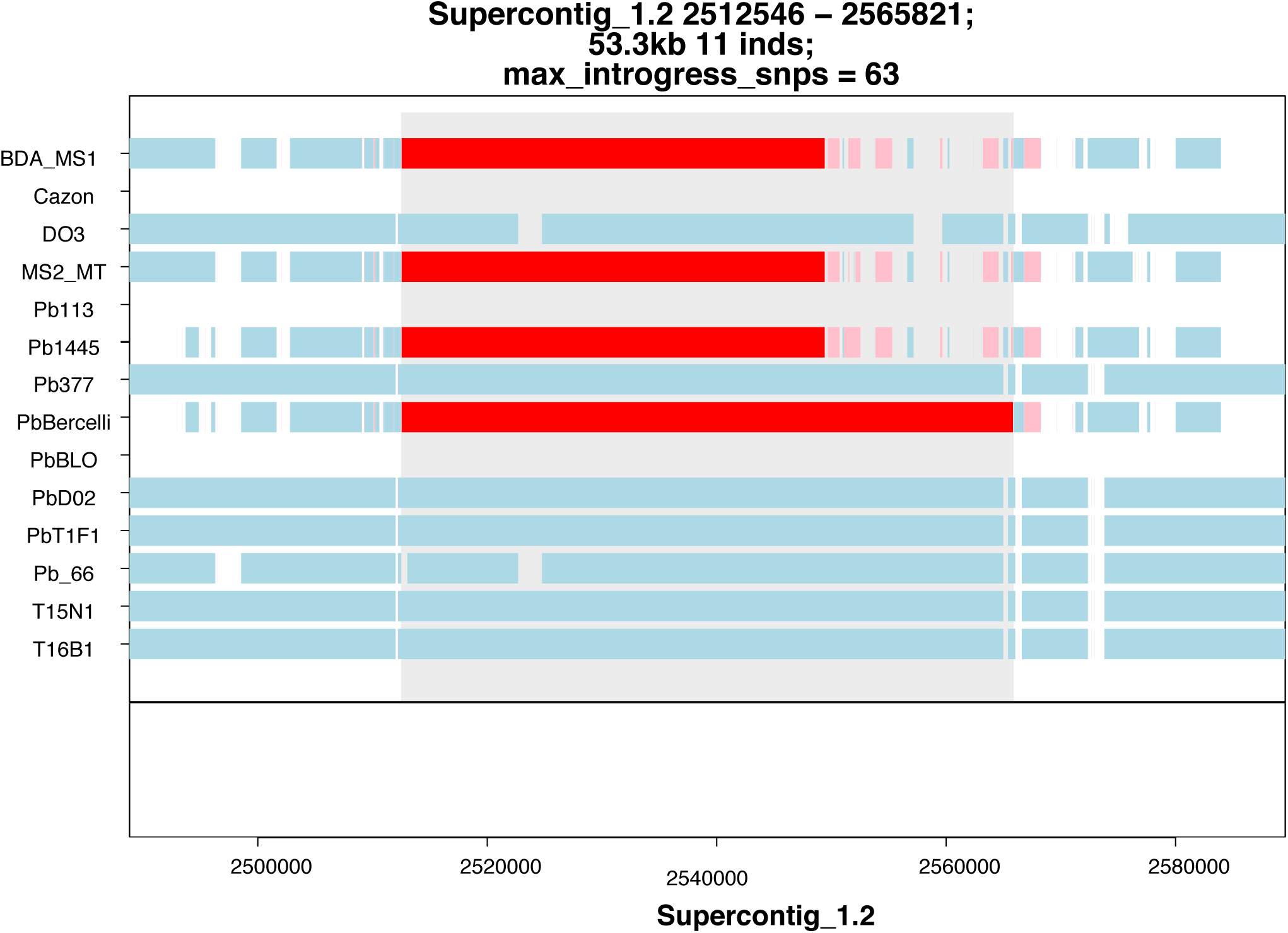
Example of an introgression from *P. restrepiensis* to *P. brasiliensis* defined by *Int-HMM*. Blue bars represent *P. brasiliensis* ancestry; red bars represent *P. restrepiensis* ancestry. Light red bars represent potential *P. restrepiensis* introgressions with low support (i.e., few markers).

**FIGURE S7.**
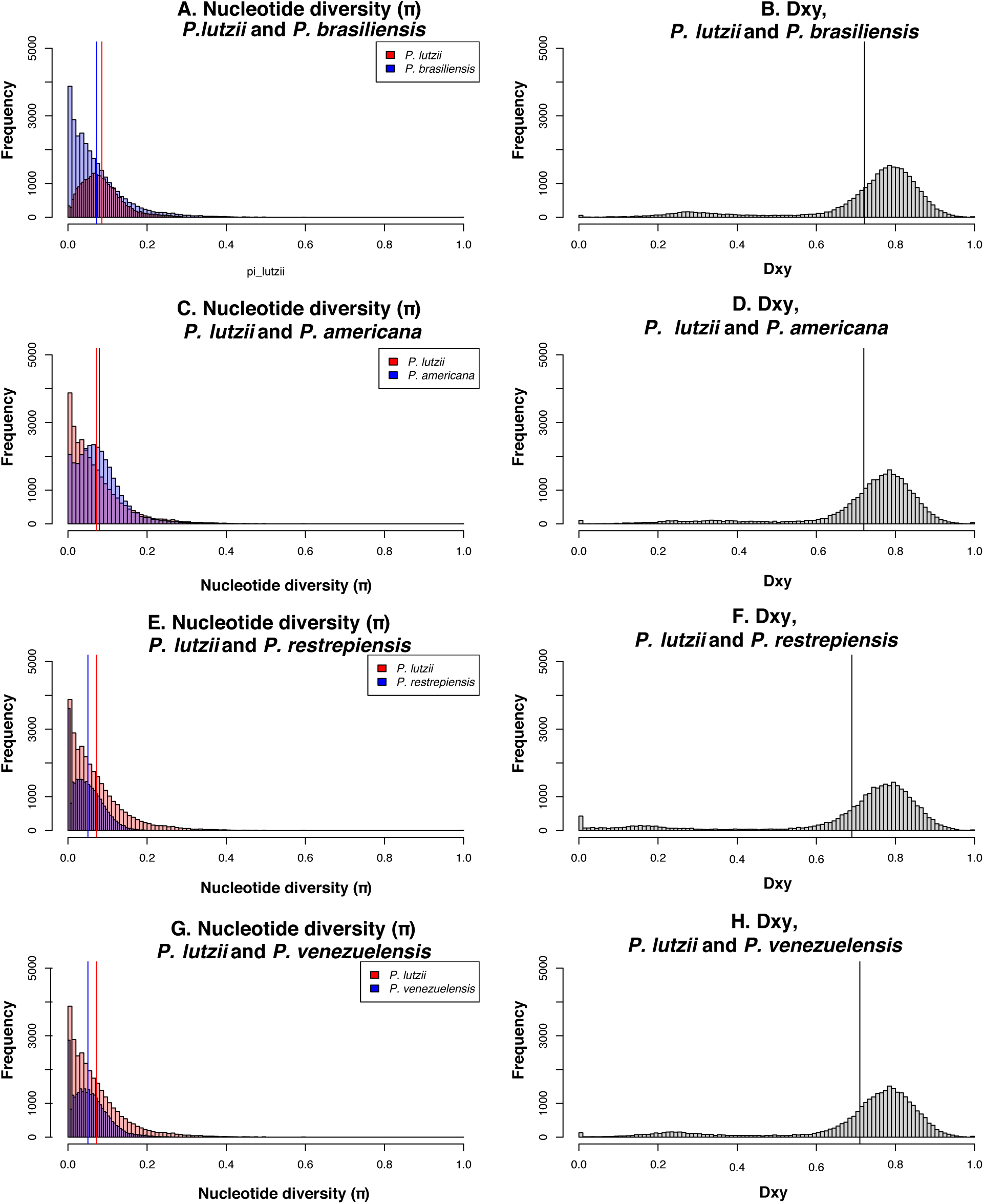
π and Dxy calculations for pairwise comparisons including *P. lutzii*. Color scheme and arrangement is similar to Figure 4. Note that Panels A and B are identical to Panels C and D in Figure 4.

**FIGURE S8.**
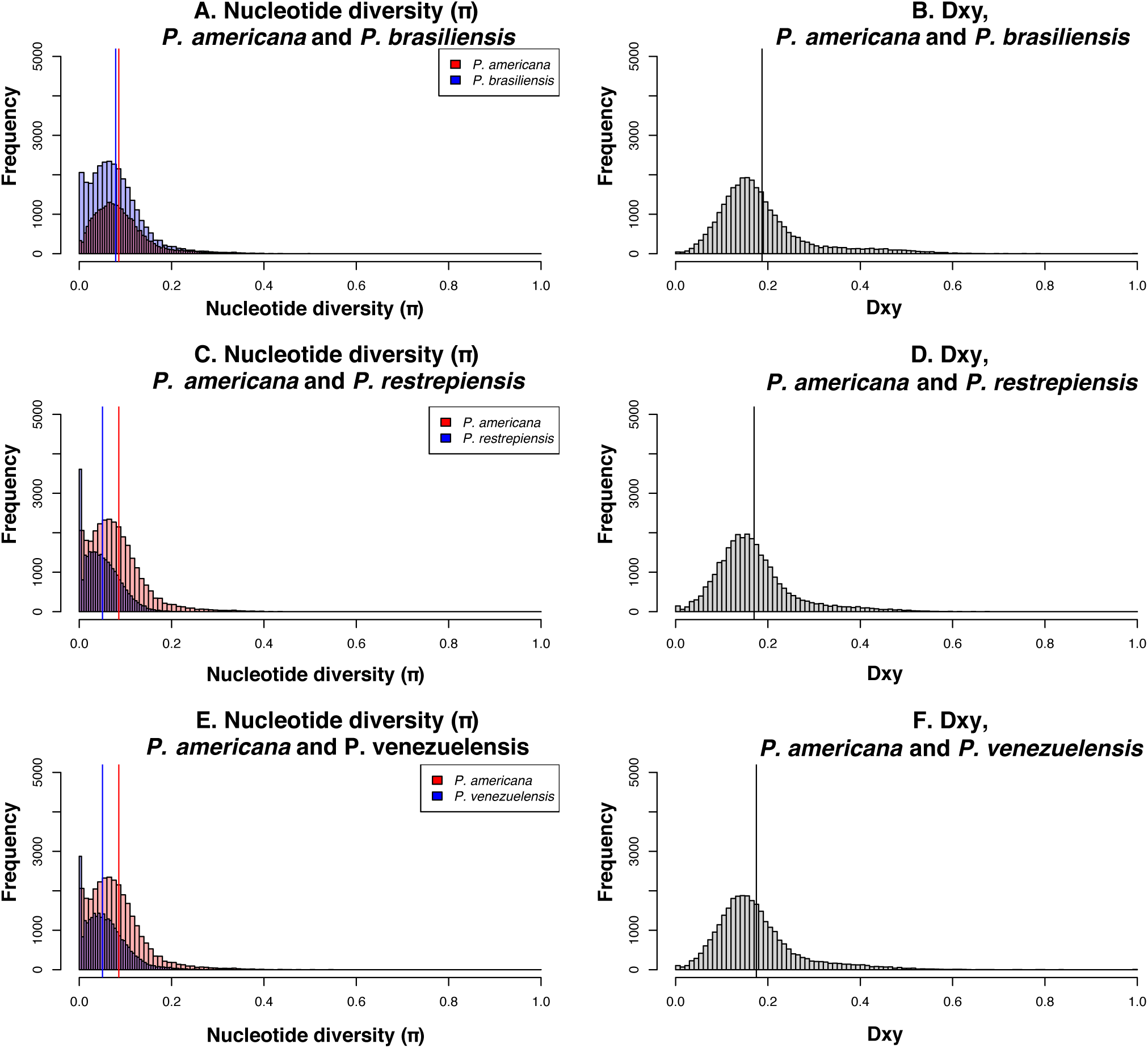
π and Dxy calculations for *P. americana*. Figure 7C-D shows the pairwise comparison between *P. americana* and *P. lutzii*.

**FIGURE S9.**
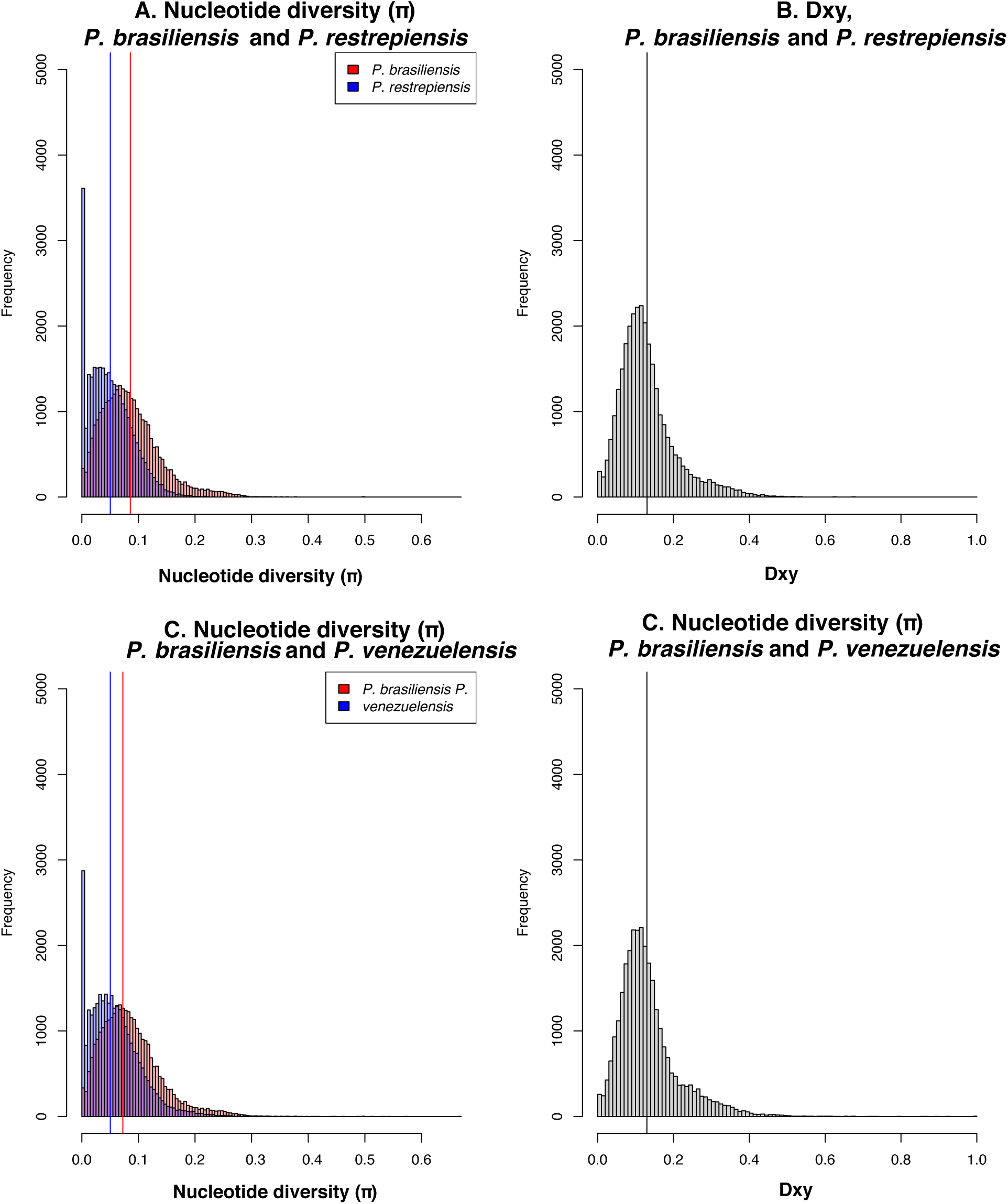
π and Dxy calculations for *P. brasiliensis*. Figure 4A-B show the comparison between *P. brasiliensis* and *P. lutzii*. Figure 8A-B shows the pairwise comparison between *P. americana* and *P. brasiliensis*.

**FIGURE S10.**
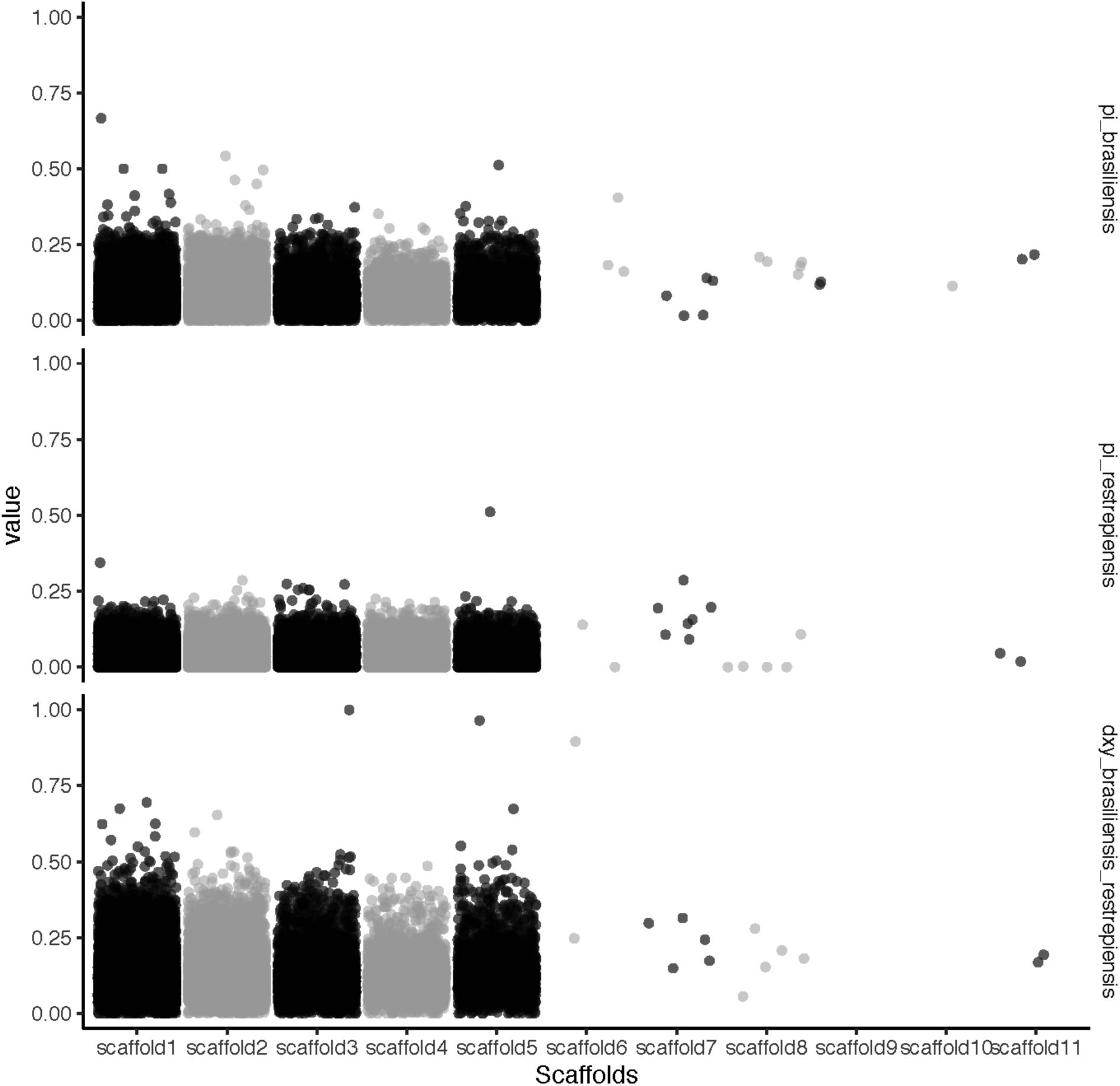
Differentiation between species of *Paracoccidioides* occurs genome-wide II. π along the whole genome within *P. brasiliensis* (upper panel) and *P. restrepiensis* (middle panel) is lower than *Dxy* between species (upper panel).

**FIGURE S11.**
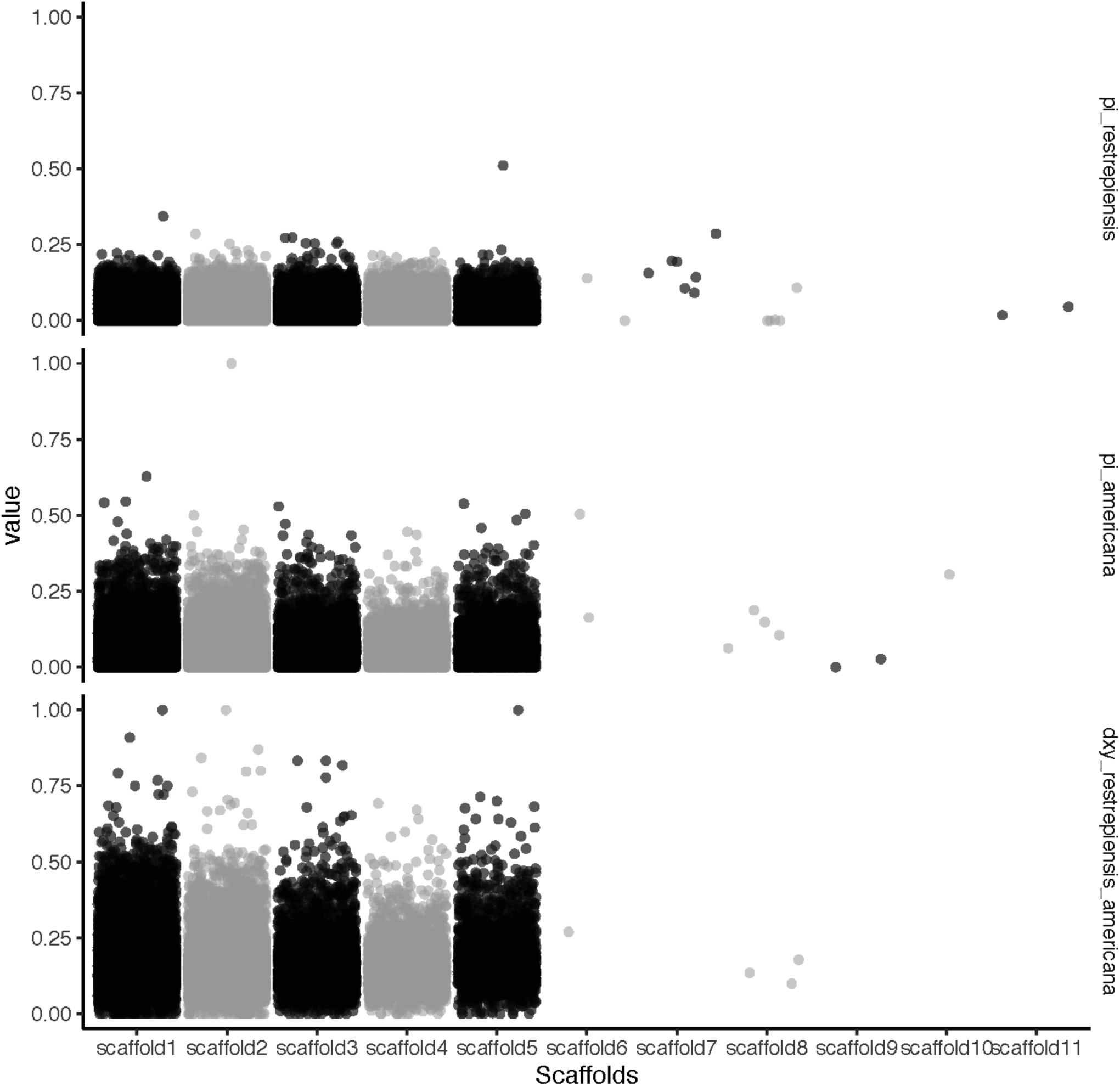
Differentiation between species of *Paracoccidioides* occurs genome-wide III. π along the whole genome within *P. restrepiensis* (upper panel) and *P. americana* (middle panel) is lower than *Dxy* between species (upper panel).

**FIGURE S12.**
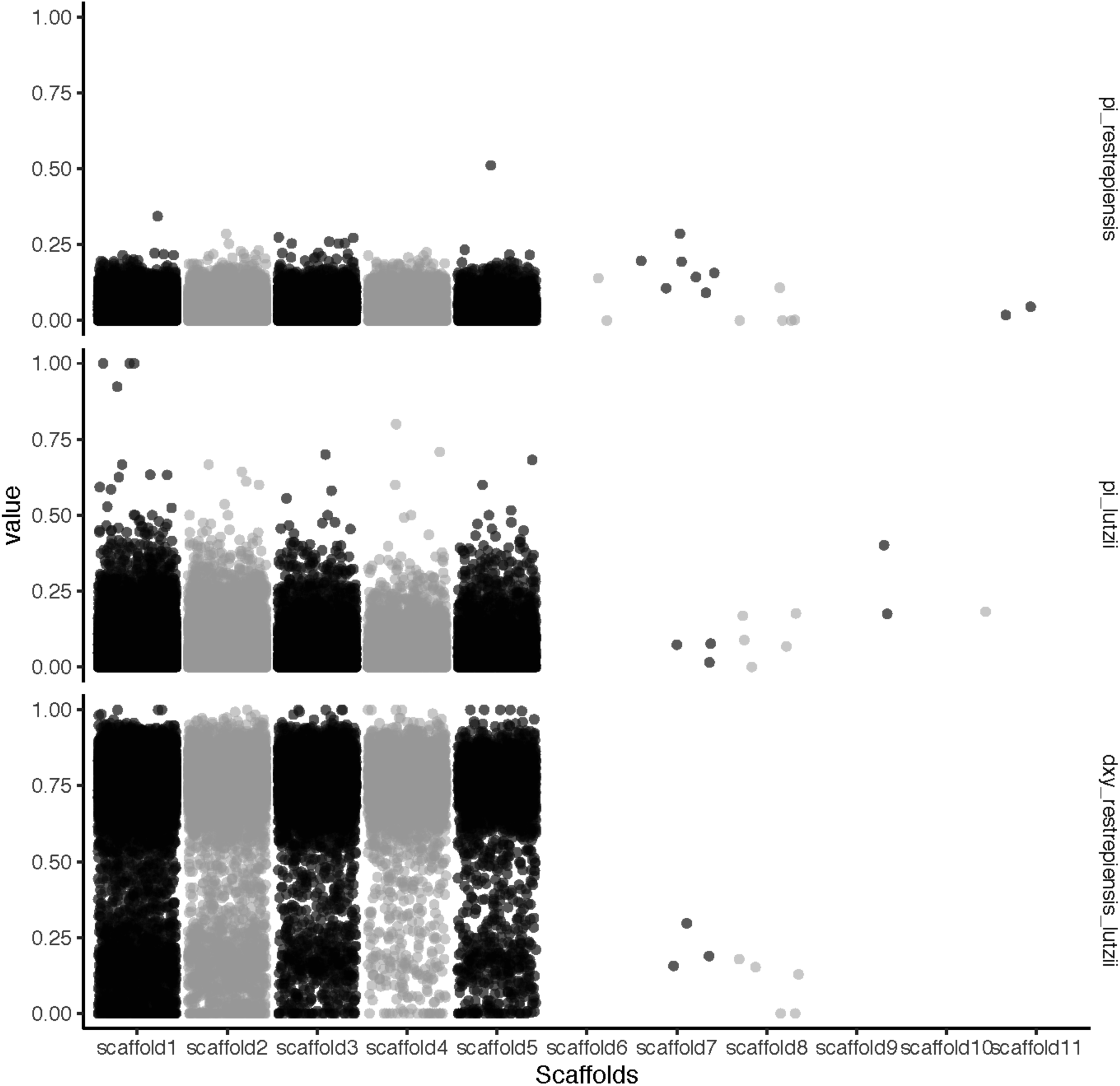
Differentiation between species of *Paracoccidioides* occurs genome-wide IV. π along the whole genome within *P. restrepiensis* (upper panel) and *P. lutzii* (middle panel) is lower than *Dxy* between species (upper panel). Other plots can be found in the data package (doi: TBD)

## REFERENCES

Almeida, A. J., D. R. Matute, J. A. Carmona, M. Martins, I. Torres, J. G. McEwen, A. Restrepo, C. Leão, P. Ludovico, and F. Rodrigues. 2007. Genome size and ploidy of *Paracoccidioides brasiliensis* reveals a haploid DNA content: Flow cytometry and GP43 sequence analysis. Fungal Genet. Biol. 44:25–31.

Ané, C. 2010. Gene tree reconciliation: new developments in Bayesian concordance analysis with BUCKy. Nat. Preced. 1:1

Ané, C., B. Larget, D. A. Baum, and S. D. Smith. 2006. Bayesian estimation of concordance among gene trees. Mol. Biol. Evol. 24: 412–426.

Arantes, T. D., R. C. Theodoro, M. de M. Teixeira, and E. Bagagli. 2017. Use of fluorescent oligonucleotide probes for differentiation between *Paracoccidioides brasiliensis* and *Paracoccidioides lutzii* in yeast and mycelial phase. Mem. Inst. Oswaldo Cruz, 112:140–145. doi: 10.1590/0074-02760160374.

Baum, D. A. 2007. Concordance trees, concordance factors, and the exploration of reticulate genealogy. Taxon, 56: 417–426.

Brummer, E., E. Castaneda, and A. Restrepo. 1993. Paracoccidioidomycosis: An update. Clinical microbiology reviews, 6: 89–117.

Chung, Y., and C. Ané. 2011. Comparing two bayesian methods for gene tree/species tree reconstruction: Simulations with incomplete lineage sorting and horizontal gene transfer. Syst. Biol., 60: 261–275. doi: 10.1093/sysbio/syr003.

Comparato Filho, O. O., F. V. Morais, T. Bhattacharjee, M. L. Castilho, and L. Raniero. 2019. Rapid identification of *Paracoccidioides lutzii* and *P. brasiliensis* using Fourier Transform Infrared spectroscopy. J. Mol. Struct., 1177: 152–159. doi: 10.1016/j.molstruc.2018.09.016.

De Almeida, J. N., G. M. B. Del Negro, R. C. Grenfell, M. S. M. Vidal, D. Y. Thomaz, D. S. Y. De Figueiredo, E. Bagagli, L. Julianod, and G. Benard. 2015. Matrix-assisted laser desorption ionization-time of flight mass spectrometry for differentiation of the dimorphic fungal species *Paracoccidioides brasiliensis* and *Paracoccidioides lutzii.* J. Clin. Microbiol., 53: 1383–1386. doi: 10.1128/JCM.02847-14.

DePristo, M. A., E. Banks, R. Poplin, K. V Garimella, J. R. Maguire, C. Hartl, A. A. Philippakis, G. del Angel, M. A. Rivas, M. Hanna, A. McKenna, T. J. Fennell, A. M. Kernytsky, A. Y. Sivachenko, K. Cibulskis, S. B. Gabriel, D. Altshuler, and M. J. Daly. 2011. A framework for variation discovery and genotyping using next-generation DNA sequencing data. Nat. Genet. 43:491–498.

Desjardins, C. A., M. D. Champion, J. W. Holder, A. Muszewska, J. Goldberg, A. M. Bailão, M. M. Brigido, M. E. da Silva Ferreira, A. M. Garcia, M. Grynberg, S. Gujja, D. I. Heiman, M. R. Henn, C. D. Kodira, H. León-Narváez, L. V. G. Longo, L. J. Ma, I. Malavazi, A. L. Matsuo, F. V. Morais, M. Pereira, S. Rodríguez-Brito, S. Sakthikumar, S. M. Salem-Izacc, S. M. Sykes, M. M. Teixeira, M. C. Vallejo, M. E. M. T. Walter, C. Yandava, S. Young, Q. Zeng, J. Zucker, M. S. Felipe, G. H. Goldman, B. J. Haas, J. G. McEwen, G. Nino-Vega, R. Puccia, G. San-Blas, C. M. A. de Soares, B. W. Birren, and C. A. Cuomo. 2011. Comparative genomic analysis of human fungal pathogens causing paracoccidioidomycosis. PLoS Genet. 7: p.e1002345.

Desjardins, C. A., C. Giamberardino, S. M. Sykes, C. H. Yu, J. L. Tenor, Y. Chen, T. Yang, A. M. Jones, S. Sun, M. R. Haverkamp, J. Heitman, A. P. Litvintseva, J. R. Perfect, and C. A. Cuomo. 2017. Population genomics and the evolution of virulence in the fungal pathogen *Cryptococcus neoformans*. Genome Res., 27: pp.1207–1219. doi: 10.1101/gr.218727.116.

Dukik, K., J. F. Muñoz, Y. Jiang, P. Feng, L. Sigler, J. B. Stielow, J. Freeke, A. Jamalian, B. Gerrits van den Ende, J. G. McEwen, O. K. Clay, I. S. Schwartz, N. P. Govender, T. G. Maphanga, C. A. Cuomo, L. F. Moreno, C. Kenyon, A. M. Borman, and S. de Hoog. 2017. Novel taxa of thermally dimorphic systemic pathogens in the Ajellomycetaceae (Onygenales). Mycoses, 60: 296–309. doi: 10.1111/myc.12601.

Durand, E. Y., N. Patterson, D. Reich, and M. Slatkin. 2011. Testing for ancient admixture between closely related populations. Mol. Biol. Evol. 28: 2239–2252.

Ellegren, H. 2004. Microsatellites: Simple sequences with complex evolution. Nature reviews genetics, 5(6), pp.435–445

Feitosa, L. D. S., P. S. Cisalpino, M. R. Machado Dos Santos, R. A. Mortara, T. F. Barros, F. V. Morais, R. Puccia, J. F. Da Silveira, and Z. P. De Camargo. 2003. Chromosomal polymorphism, syntenic relationships, and ploidy in the pathogenic fungus *Paracoccidioides brasiliensis*. Fungal Genet. Biol., 39: 60–69. doi: 10.1016/S1087-1845(03)00003-3.

Gao, Z., M. Przeworski, and G. Sella. 2015. Footprints of ancient-balanced polymorphisms in genetic variation data from closely related species. Evolution 69:431–446.

Giraud, T., and S. Gourbière. 2012. The tempo and modes of evolution of reproductive isolation in fungi. Heredity (Edinb). 109:204–214.

Green, R. E., J. Krause, A. W. Briggs, T. Maricic, U. Stenzel, M. Kircher, N. Patterson, H. Li, W. Zhai, M. H.-Y. Fritz, N. F. Hansen, E. Y. Durand, A.-S. Malaspinas, J. D. Jensen, T. Marques-Bonet, C. Alkan, K. Prüfer, M. Meyer, H. A. Burbano, J. M. Good, R. Schultz, A. Aximu-Petri, A. Butthof, B. Höber, B. Höffner, M. Siegemund, A. Weihmann, C. Nusbaum, E. S. Lander, C. Russ, N. Novod, J. Affourtit, M. Egholm, C. Verna, P. Rudan, D. Brajkovic, Ž. Kucan, I. Gušic, V. B. Doronichev, L. V Golovanova, C. Lalueza-Fox, M. de la Rasilla, J. Fortea, A. Rosas, R. W. Schmitz, P. L. F. Johnson, E. E. Eichler, D. Falush, E. Birney, J. C. Mullikin, M. Slatkin, R. Nielsen, J. Kelso, M. Lachmann, D. Reich, and S. Pääbo. 2010. A Draft Sequence of the Neandertal Genome. Science (80-.). 328:710–722.

Hamlin, J. A. P., M. S. Hibbins, and L. C. Moyle. 2020. Assessing biological factors affecting postspeciation introgression. Evol. Lett., 4: 137–154. doi: 10.1002/evl3.159.

Hendry, A. P., D. I. Bolnick, D. Berner, and C. L. Peichel. 2009. Along the speciation continuum in sticklebacks. J. Fish Biol., 75: 2000–2036. doi: 10.1111/j.1095-8649.2009.02419.x.

Hhna, S., B. Larget, L. Liu, M. A. Suchard, and J. P. Huelsenbeck. 2012. MrBayes 3.2: efficient Bayesian phylogenetic inference and model choice across a large model space. Syst Biol. 61: 539–542.

Hothorn, T., K. Hornik, M. A. Van De Wiel, and A. Zeileis. 2008. Implementing a class of permutation pests: the coin package. Journal of statistical software, 28: 1–23.

Hudson, R. R., and J. A. Coyne. 2002. Mathematical consequences of the genealogical species concept. Evolution, 56(8), pp.1557-1565. dx.doi.org 56:1557.

Hughes, K. W., R. H. Petersen, and E. B. Lickey. 2009. Using heterozygosity to estimate a percentage DNA sequence similarity for environmental species’ delimitation across basidiomycete fungi. New Phytol. 182: 795–798.

Jarne, P., and P. J. L. Lagoda. 1996. Microsatellites, from molecules to populations and back. Trends Ecol. Evol, 11: 424–429.

Joly, S., P. A. McLenachan, and P. J. Lockhart. 2015. A statistical approach for distinguishing hybridization and incomplete lineage sorting. Am. Nat. 174:E54--E70.

Juric, I., S. Aeschbacher, and G. Coop. 2016. The strength of selection against Neanderthal introgression. PLoS Genet. 12:e1006340.

Katoh, K., and D. M. Standley. 2013. MAFFT multiple sequence alignment software version 7: improvements in performance and usability. Mol. Biol. Evol. 30: 772–780.

Kurokawa, C. S., C. R. Lopes, M. F. Sugizaki, E. E. Kuramae, M. F. Franco, and M. T. S. Peraçoli. 2005. Virulence profile of ten *Paracoccidioides brasiliensis* isolates. Association with morphologic and genetic patterns. Rev. Inst. Med. Trop. Sao Paulo, 47: 257–262. doi: 10.1590/s0036-46652005000500004.

Larget, B. R., S. K. Kotha, C. N. Dewey, and C. Ané. 2010. BUCKy: Gene tree/species tree reconciliation with Bayesian concordance analysis. Bioinformatics, 26: 2910–2911. doi: 10.1093/bioinformatics/btq539.

Lenhard-Vidal, A., J. P. Assolini, M. A. Ono, C. S. O. Bredt, A. Sano, and E. N. Itano. 2013. *Paracoccidioides brasiliensis* and *P. lutzii* antigens elicit different serum IgG responses in chronic paracoccidioidomycosis. Mycopathologia, 176: 345–352.doi: 10.1007/s11046-013-9698-0.

Li, H. 2013. Aligning sequence reads, clone sequences and assembly contigs with BWA-MEM. arXiv Prepr. arXiv 00:3.

Liepelt, S., V. Kuhlenkamp, M. Anzidei, G. G. Vendramin, and B. Ziegenhagen. 2001. Pitfalls in determining size homoplasy of microsatellite loci. Mol. Ecol. Notes, 1: 332–335. doi: 10.1046/j.1471-8278.2001.00097.x.

Malinsky, M. 2019. Dsuite - fast D-statistics and related admixture evidence from VCF files. bioRxiv, doi: 10.1101/634477.

Martin, S. H., K. K. Dasmahapatra, N. J. Nadeau, C. Salazar, J. R. Walters, F. Simpson, M. Blaxter, A. Manica, J. Mallet, and C. D. Jiggins. 2013. Genome-wide evidence for speciation with gene flow in *Heliconius* butterflies. Genome Res. 23:1817–1828.

Martin, S. H., J. W. Davey, and C. D. Jiggins. 2014. Evaluating the use of ABBA/BABA statistics to locate introgressed loci. Mol. Biol. Evol. 32(1), pp.244–257.

Martin, S. H., J. W. Davey, C. Salazar, and C. D. Jiggins. 2019. Recombination rate variation shapes barriers to introgression across butterfly genomes. PLoS Biol., 17: e2006288. doi: 10.1371/journal.pbio.2006288.

Matute, D. R., I. A. Butler, D. A. Turissini, and J. A. Coyne. 2010. A test of the snowball theory for the rate of evolution of hybrid incompatibilities. Science (80-.). 329:1518–1521.

Matute, D. R., J. G. McEwen, R. Puccia, B. A. Montes, G. San-Blas, E. Bagagli, J. T. Rauscher, A. Restrepo, F. Morais, G. Niño-Vega, and J. W. Taylor. 2006. Cryptic speciation and recombination in the fungus *Paracoccidioides brasiliensis* as revealed by gene genealogies. Mol. Biol. Evol., 23: 65–73. doi: 10.1093/molbev/msj008.

Matute, D. R., and V. E. Sepúlveda. 2019. Fungal species boundaries in the genomics era. Fungal Genetics and Biology, 131: 103249.

Maxwell, C. S., K. Mattox, D. A. Turissini, M. M. Teixeira, B. M. Barker, and D. R. Matute. 2018a. Gene exchange between two divergent species of the fungal human pathogen, *Coccidioides*. Evolution, 73: 42–58. doi: 10.1111/evo.13643.

Maxwell, C. S., V. E. Sepulveda, D. A. Turissini, W. E. Goldman, and D. R. Matute. 2018b. Recent admixture between species of the fungal pathogen *Histoplasma*. Evol. Lett. 2:210–220.

McKenna, A., M. Hanna, E. Banks, A. Sivachenko, K. Cibulskis, A. Kernytsky, K. Garimella, D. Altshuler, S. Gabriel, M. Daly, and M. A. DePristo. 2010. The Genome Analysis Toolkit: a MapReduce framework for analyzing next-generation DNA sequencing data. Genome Res. 20:1297–1303.

Mendes, F. K., and M. W. Hahn. 2018. Why concatenation fails near the anomaly zone. Syst. Biol., 67: 158–169. doi: 10.1093/sysbio/syx063.

Morais, F. V., T. F. Barros, M. K. Fukada, P. S. Cisalpino, and R. Puccia. 2000. Polymorphism in the gene coding for the immunodominant antigen gp43 from the pathogenic fungus *Paracoccidioides brasiliensis*. J. Clin. Microbiol. 38: 3960–3966.

Moyle, L. C., and T. Nakazato. 2010. Hybrid incompatibility snowballs between *Solanum* species. Science (80-.). 329:1521–1523.

Muirhead, C. A., and D. C. Presgraves. 2016. Hybrid Incompatibilities, local adaptation, and the genomic distribution of natural introgression between species. Am. Nat. 187:249–261.

Muñoz JF, Farrer RA, D. C., M. E. Gallo JE, Sykes S, Sakthikumar S, T. Whiston EA, Bagagli E, Soares CMA, M. J. MDM, Taylor JW, Clay OK, C. Cuomo, and CA. 2016. Genome diversity, recombination and virulence across the major lineages of *Paracoccidioides*. mSphere, 1: e00213–16. doi: 10.1128/mSphere.00213-16.

Niño-Vega, G. A., A. M. Calcagno, G. San-Blas, F. San-Blas, G. W. Gooday, and N. A. R. Gow. 2000. RFLP analysis reveals marked geographical isolation between strains of *Paracoccidioides brasiliensis.* Med. Mycol., 38: 437–441. doi: 10.1080/mmy.38.6.437.441.

Pereira de Souza, S., V. Magalhães Jorge, and M. Orzechowski Xavier. 2014. Paracoccidioidomycosis in southern Rio Grande do Sul: A retrospective study of histopathologically diagnosed cases. Brazilian J. Microbiol. 45: 243–247.

Pryszcz L. P., Gabaldón T. (2016). Redundans: an assembly pipeline for highly heterozygous genomes. Nucleic Acids Res. 44:e113. 10.1093/nar/gkw294

Queiroz, K. De, and K. De Queiroz. 2007. Species concepts and species delimitation. Syst. Biol., 56: 879–886. doi: 10.1080/10635150701701083.

R Core Team. 2016. R Development Core Team.

Restrepo, A. M., A. M. Tobon Orozco, B. L. Gomez, and G. Benard. 2015. Paracoccidioidomycosis. P. in Diagnosis and Treatment of Fungal Infections.

Restrepo, A., J. G. McEwen, and E. Castañeda. 2001. The habitat of *Paracoccidioides brasiliensis*: how far from solving the riddle?. Med. Mycol., 39: 233–241. doi: 10.1080/mmy.39.3.233.241.

Restrepo, A., and A. M. Tobón. 2010. Paracoccidioides brasiliensis. P. in Mandell, Douglas, and Bennett’s Principles and Practice of Infectious Diseases.

Restrepo M, A. 1985. The ecology of *Paracoccidioides brasiliensis*: A puzzle still unsolved. Sabouraudia: Journal of Medical and Veterinary Mycology, 23: 323–334.

Richini-Pereira, V. B., S. D. M. G. Bosco, J. Griese, R. C. Theodoro, S. A. D. G. Macoris, R. J. Da Silva, L. Barrozo, P. M. E. S. Tavares, R. M. Zancopé-Oliveira, and E. Bagagli. 2008. Molecular detection of *Paracoccidioides brasiliensis* in road-killed wild animals. Med. Mycol., 46: 35–40. doi: 10.1080/13693780701553002.

Richini-Pereira, V., S. Bosco, R. Theodoro, L. Barrozo, S. Pedrini, P. Rosa, and E. Bagagli. 2009. Importance of xenarthrans in the eco-epidemiology of *Paracoccidioides brasiliensis*. BMC Res. Notes, 2: 228. doi: 10.1186/1756-0500-2-228.

Roberto, T. N., A. M. Rodrigues, R. C. Hahn, and Z. P. De Camargo. 2016. Identifying *Paracoccidioides* phylogenetic species by PCR-RFLP of the alpha-tubulin gene. Med. Mycol., 54: 240–247. doi: 10.1093/mmy/myv083.

Ronquist, F., and J. P. Huelsenbeck. 2003. MrBayes 3: Bayesian phylogenetic inference under mixed models. Bioinformatics 19:1572–1574.

Ronquist, F., M. Teslenko, P. van der Mark, D. L. Ayres, A. Darling, S. Hohna, B. Larget, L. Liu, M. A. Suchard and J. P. Huelsenbeck. 2012. MrBayes 3.2: efficient Bayesian phylogenetic inference and model choice across a large model space. Syst Biol 61: 539-542. doi. 10.1093/sysbio/sys029

Roux, C., C. Fraïsse, J. Romiguier, Y. Anciaux, N. Galtier, and N. Bierne. 2016. Shedding Light on the Grey Zone of Speciation along a Continuum of Genomic Divergence. PLoS Biol., 14: e2000234. doi: 10.1371/journal.pbio.2000234.

Salgado-Salazar, C., L. R. Jones, Á. Restrepo, and J. G. McEwen. 2010. The human fungal pathogen *Paracoccidioides brasiliensis* (Onygenales: Ajellomycetaceae) is a complex of two species: Phylogenetic evidence from five mitochondrial markers. Cladistics, 26: 613–624. doi: 10.1111/j.1096-0031.2010.00307.x.

Sankararaman, S., S. Mallick, M. Dannemann, K. Prüfer, J. Kelso, S. Pääbo, N. Patterson, and D. Reich. 2014. The genomic landscape of Neanderthal ancestry in present-day humans. Nature 507:354–357.

Schumer, M., C. Xu, D. L. Powell, A. Durvasula, L. Skov, C. Holland, J. C. Blazier, S. Sankararaman, P. Andolfatto, G. G. Rosenthal, and M. Przeworski. 2018. Natural selection interacts with recombination to shape the evolution of hybrid genomes. Science (80-.)., 360: 656–660. doi: 10.1126/science.aar3684.

Sepúlveda, V. E., R. Márquez, D. A. Turissini, W. E. Goldman, and D. R. Matute. 2017. Genome sequences reveal cryptic speciation in the human pathogen *Histoplasma capsulatum*. MBio 8:e01339–17.

Silva Vergara, M. L., and R. Martinez. 1998. Role of the armadillo *Dasypus novemcinctus* in the epidemiology of paracoccidioidomycosis. Mycopathologia, 144: 131–133. doi: 10.1023/A:1007034215003.

Simão, F. A., R. M. Waterhouse, P. Ioannidis, E. V. Kriventseva, and E. M. Zdobnov. 2015. BUSCO: Assessing genome assembly and annotation completeness with single-copy orthologs. Bioinformatics, 31: 3210–3212. doi: 10.1093/bioinformatics/btv351.

Siqueira, I. M., C. L. F. Fraga, A. C. Amaral, A. C. O. Souza, M. S. Jerônimo, J. R. Correa, K. G. Magalhães, C. A. Inácio, A. M. Ribeiro, P. H. Burguel, M. S. Felipe, A. H. Tavares, and A. L. Bocca. 2016. Distinct patterns of yeast cell morphology and host responses induced by representative strains of *Paracoccidioides brasiliensis* (Pb18) and *Paracoccidioides lutzii* (Pb01). Med. Mycol., 54: 177–188. doi: 10.1093/mmy/myv072.

Stamatakis, A. 2006. RAxML-VI-HPC: maximum likelihood-based phylogenetic analyses with thousands of taxa and mixed models. Bioinformatics 22:2688–2690.

Taylor, J. W., D. J. Jacobson, S. Kroken, T. Kasuga, D. M. Geiser, D. S. Hibbett, and M. C. Fisher. 2000. Phylogenetic species recognition and species concepts in fungi. Fungal Genet. Biol., 31: 21–32. doi: 10.1006/fgbi.2000.1228.

Teixeira, M., R. C. Theodoro, F. F. M. De Oliveira, G. C. Machado, R. C. Hahn, E. Bagagli, G. San-Blas, and M. S. S. Felipe. 2015. *Paracoccidioides lutzii s*p. nov.: Biological and clinical implications. Med. Mycol., 52: 19–28.

Teixeira, M. de M., M. E. Cattana, D. R. Matute, J. F. Muñoz, A. Arechavala, K. Isbell, Schipper, G. Santiso, F. Tracogna, M. de los Á. Sosa, N. Cech, P. Alvarado, L. Barreto, Y. Chacón, J. Ortellado, C. M. de Lima, M. R. Chang, G. Niño-Vega, M. A. Yasuda, M. S. S. Felipe, R. Negroni, C. A. Cuomo, B. Barker, and G. Giusiano. 2020. Genomic diversity of the human pathogen *Paracoccidioides* across the South American continent. Fungal Genet. Biol., doi: 10.1016/j.fgb.2020.103395.

Teixeira, M. de M., R. C. Theodoro, L. da S. Derengowski, A. M. Nicola, E. Bagagli, and M. S. Felipe. 2013. Molecular and morphological data support the existence of a sexual cycle in species of the genus *Paracoccidioides*. Eukaryot. Cell, 12: 380–389. doi: 10.1128/EC.05052-11.

Teixeira, M. M., R. C. Theodoro, M. J. A. de Carvalho, L. Fernandes, H. C. Paes, R. C. Hahn, L. Mendoza, E. Bagagli, G. San-Blas, and M. S. S. Felipe. 2009. Phylogenetic analysis reveals a high level of speciation in the *Paracoccidioides* genus. Mol. Phylogenet. Evol., 52: 273–283. doi: 10.1016/j.ympev.2009.04.005.

Teixeira, M. M., R. C. Theodoro, G. Nino-Vega, E. Bagagli, and M. S. S. Felipe. 2014. *Paracoccidioides* species complex: ecology, phylogeny, sexual reproduction, and virulence. PLoS Pathog., 10: e1004397. doi: 10.1371/journal.ppat.1004397.

Turissini, D. A., O. M. Gomez, M. M. Teixeira, J. G. McEwen, and D. R. Matute. 2017. Species boundaries in the human pathogen *Paracoccidioides*. Fungal Genet. Biol. 106:9–25. doi: 10.1016/j.fgb.2017.05.007.

Turissini, D. A., and D. R. Matute. 2017. Fine scale mapping of genomic introgressions within the *Drosophila yakuba clade*. PLoS Genet. 13:e1006971.

Wang, R. J., M. A. White, and B. A. Payseur. 2015. The pace of hybrid incompatibility evolution in house mice. Genetics 201:229–242.

Waterhouse, R. M., M. Seppey, F. A. Simao, M. Manni, P. Ioannidis, G. Klioutchnikov, E. V. Kriventseva, and E. M. Zdobnov. 2018. BUSCO applications from quality assessments to gene prediction and phylogenomics. Mol. Biol. Evol., 35: 543–548. doi: 10.1093/molbev/msx319.

